# Structural basis of Omicron immune evasion: A comparative computational study of Spike protein-Antibody interaction

**DOI:** 10.1101/2022.03.15.484421

**Authors:** Darshan Contractor, Christoph Globisch, Shiv Swaroop, Alok Jain

## Abstract

The COVID-19 pandemic has caused more than 424 million infections and 5.9 million deaths so far. The vaccines used against SARS-COV-2 by now have been able to develop some neutralising antibodies in the vaccinated human population and slow down the infection rate. The effectiveness of the vaccines has been challenged by the emergence of the new strains with numerous mutations in the spike (S) protein of SARS-CoV-2. Since S protein is the major immunogenic protein of the virus and also contains Receptor Binding Domain (RBD) that interacts with the human Angiotensin-Converting Enzyme 2 (ACE2) receptors, any mutations in this region should affect the neutralisation potential of the antibodies leading to the immune evasion. Several variants of concern (VOC) of the virus have emerged so far. Among them, the most critical are Delta (B.1.617.2), and recently reported Omicron (B. 1.1.529) which have acquired a lot of mutations in the spike protein. We have mapped those mutations on the modelled RBD and evaluated the binding affinities of various human antibodies with it. Docking and molecular dynamics simulation studies have been used to explore the effect of the mutations on the structure of the RBD and the RBD-antibody interaction. The analysis shows that the mutations mostly at the interface of a nearby region lower the binding affinity of the antibody by ten to forty per cent, with a downfall in the number of interactions formed as a whole and therefore, it implies the generation of immune escape variants. Notable mutations and their effect was characterised by performing various analyses that explain the structural basis of antibody efficacy in Delta and a compromised neutralisation effect for the Omicron variant. Our results pave the way for robust vaccine design that can be effective for many variants.

**Graphical Abstract:** 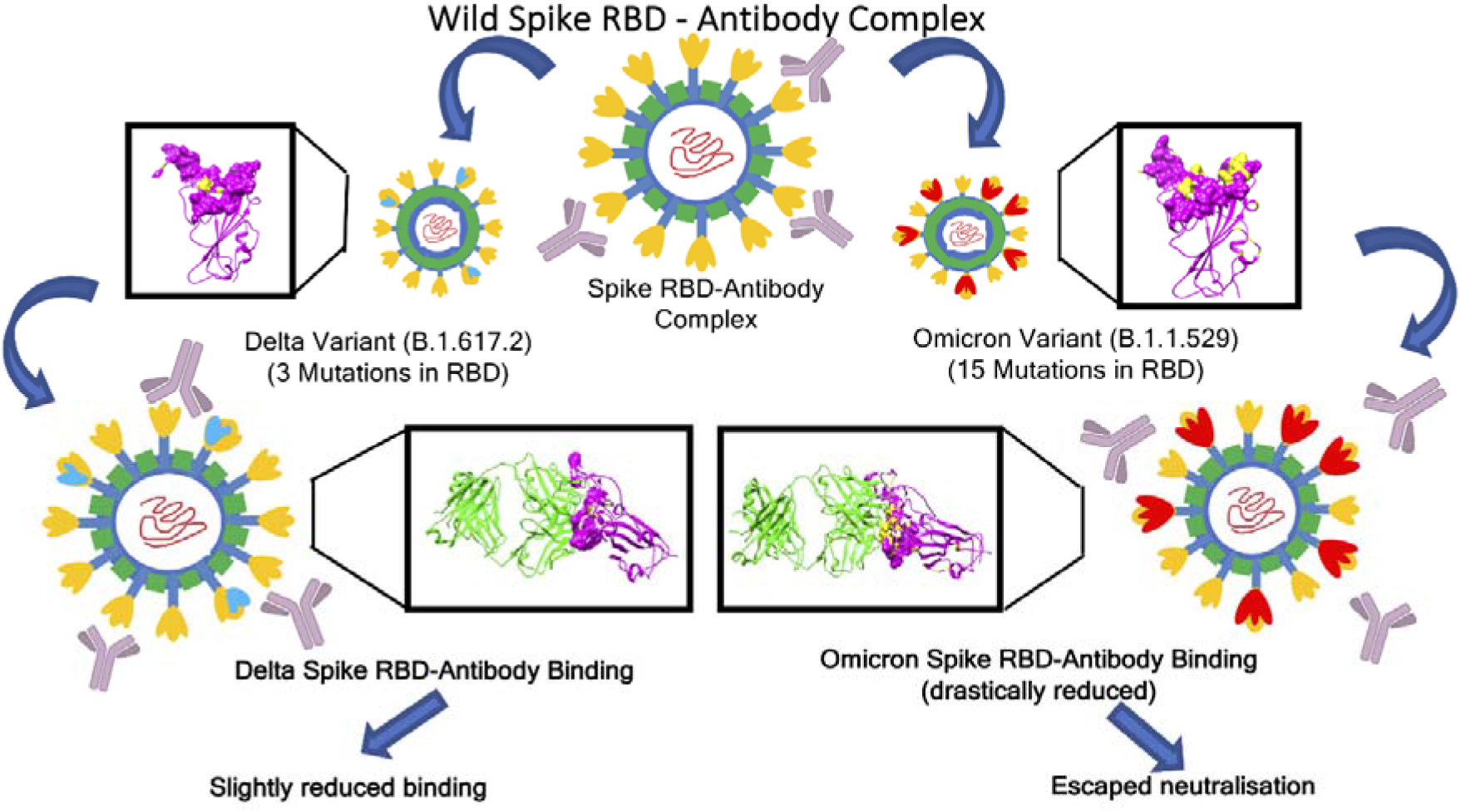

**Synopsis:** The research study utilises comparative docking and MD simulations analyses to illustrate how mutations in delta and omicron variants affect the binding of antibodies to the spike receptor binding domain (RBD) of SARS CoV-2.

## Introduction

Human Coronavirus disease 2019 (COVID-19) is an extremely infectious disease which is caused by SARS coronavirus 2 (SARS-CoV-2). The disease was first reported in Wuhan, China in mid-December 2019 and has spread globally after that, resulting in the ongoing pandemic. Symptoms vary from mild in majority of the cases to severe like pneumonia and multi-organ failure in few^1,2^.

The genome of SARS-CoV-2 is a single-stranded positive-sense RNA that codes for 10 genes and 26 proteins. The surface spike protein (S) is not only important for the viral entry but also is the main immunogenic protein in the virus^3^. The S protein is known to be cleaved into an amino-terminal S1 subunit which is involved in virus–host cell receptor binding, and a carboxyl-terminal S2 subunit that is responsible for a virus–host membrane fusion. The S1 subunit contains two domains, an N-terminal (NTD) and a C-terminal (CTD) domain where the latter called receptor binding domain (RBD) as it is involved in receptor binding. The RBD of S protein is a promising target for molecules which can bind to RBD cause viral entry inhibition^4,5^.

Vaccines are used as a main prophylactic measure against any type of infection. They mimic the natural infection in terms of activating the host immune response, including antibody generation, to develop an immunity against that pathogen. The use of vaccines against SARS-CoV-2 has been a major success in reducing the pace of COVID-19 pandemic. Till now, more than 100 vaccines of different types have been developed, and around 26 vaccines have undergone phase III clinical trials, as per WHO^6^.

Till date, there are many vaccines available in the market across the world. Some of the main vaccines are: mRNA vaccines (BNT16b2, mRNA-1273, CVnCoV), viral vector vaccines (AZD1222, Sputnik V, Sputnik V Light, Ad5-nCoV (Convidecia), Ad26. (COV2.S), inactivated vaccines (NVX-COV2373, CoronaVac, BBIBP-CorV, Wuhan Sinopharm inactivated vaccine, Covaxin, QazVac, KoviVac, COVIran Barekat), and protein-based vaccines (EpiVacCorona, ZF2001, Abdala)^7^. One major challenge associated with the success of vaccines is the fast mutation rate of SARS-CoV-2. Since the initial outbreak, several variants have emerged which include the variants of concerns (VOCs) like Alpha (B.1.1.7), Beta (B.1.351), Gamma (P.1) and Delta (B.1.617.2) lineages. The mutations in these VOCs are the cause for several infection waves and increased transmission or mortality of COVID-19^8–12^. The specific mutations in that region of the spike protein which is the target of the antibodies leads to escaping the immune response when compared to the original Wuhan strain or D614G variant^13–18^.

Till November 2021, the delta variant of the SARS-CoV-2 was the main variant of concern as it could spread over 163 nations by August 2021^19^. On 26^th^ November 2021, a new variant B. 1.1.529, commonly called Omicron, was declared as a VOC by WHO. This mutant was discovered in Botswana and South Africa in mid-November, 2021. As per the reports so far, Omicron has several mutations which increased its transmissibility manifold as compared to the Delta variant. The spike protein in Omicron has thirty mutations, fifteen of which are present in the receptor-binding domain^20, 21^. The original Wuhan strain against which the initial vaccines were developed may not be effective against the new variants due the continuous mutations in the spike protein.

There are several experimental and computational studies which clearly show the cause of a high transmission of the Omicron variant and its immune evasion^22–25^. The increase in the binding of the SARS-CoV-2 receptor ACE2 with the RBD is the main cause of the enhanced transmission of the Omicron variant. The RBD-ACE2 receptor interaction is very well characterised in the case of VOCs that emerged before Omicron and has been evidenced as the major cause of higher transmission in the Delta variant. Kumar *et al*.^23^ performed computational studies to assess the effect of spike protein mutations in the Delta and Omicron variants. It was found that the Omicron variant showed higher binding affinity with the human ACE-2 receptor as compared to Delta variant. In another study, it was shown that the mutations in the Omicron RBD yields more contacts, hydrogen bonds and buried surface area at the interface of spike RBD and ACE-2 receptor, as compared to the original strain^24^. Rath *et al.* ^22^ reported 2.5 times stronger binding between the mutated residues and ACE-2 receptor together with a much relaxed dynamics of the complex, as compared to the wild type.

In this study we focus on finding the molecular basis of immune evasion by the Omicron variant, as it is already known to infect the previously infected as well as vaccinated population. Cao *et al*. ^26^ determined the escape mutations in the RBD against a panel of 247 anti-RBD Nabs and found that several single Omicron mutations lead to impaired Nabs of various epitopes. In another study, an infectious Omicron virus isolated in Belgium was tested for its sensitivity to the antibodies in the sera of 115 people which were either vaccinated or recovered from COVID infection, and against nine clinically approved monoclonal antibodies (mAbs). Omicron showed partial or total resistance to neutralisation by all mAbs. Similar results were observed for the sera from vaccinated individuals and convalescent sera collected 5 to 6 months after the vaccination or infection of individuals^27^. In a related study, Liu *et al*. ^24^ reported diminished neutralisation of the Omicron by the convalescent sera and sera from the vaccinated individuals. Similar results were found for 17 out of the 19 mAbs specific to all known epitope clusters on the spike protein^27^.

In this article, we used computational tools like molecular docking, MD simulations, free energy calculations and thorough structure analysis to assess the effect of such mutations in the Delta and the Omicron variants on the structure of RBD and its binding with the major neutralising antibodies generated against SARS-COV-2. These antibodies include CR3022^28^, S230^29^, CC12.1^30^, REGN10987^31^ and S309^32^. The results show that the mutations in the RBD and few at other positions in the spike protein lowers the binding affinity between the RBD and the various antibodies. We also looked into the original interactions which are hampered and the new interactions resulting out of these mutations. Such studies are useful not only to understand the cause of immune evasion by these VOCs at a molecular level, but also to predict the immune evasion ability of the upcoming variants and act accordingly for the preparedness against those variants.

## Results and Discussions

### Characterisation of neutralising antibodies via Molecular Docking

Any protein-protein or protein-ligand docking is represented in the form of ‘best fit’ orientation between them which is the principle of molecular docking. The binding affinity of the various neutralising antibodies to that of the receptor binding domain of the spike protein was evaluated by molecular docking analysis.

The RBD of the spike proteins from three variants of SARS-CoV-2, the wild type, the delta (B.1.617.2) and the omicron (B.1.1.529) were docked with the various antibodies as mentioned in the materials and method section, and the interactions among the residues were analysed. The protein-protein docking analysis was accomplished using the web based tool - ClusPro 2.0^33^ and the results were compared and interpreted. In the obtained structures the antibodies bind to different epitope sites of the viral RBD forming various interactions among the spike and the antibody residues. Relative binding locations of five antibodies considered in our study are shown in Figure-1A/B as observed in the crystal structures. It is evident that the epitopes of CC12.1, REGN10987, CR3022 and S230 are in the close proximity of mutant residues (Figure 1) of Omicron and Delta strains respectively. This Proximity of mutant residues to antibody binding sites may affect their neutralising efficiency as compared to the wild type strain. However, to our surprise no significant changes in the docking scores were observed (Table S1), which was in contrast to the published data that clearly showed a significant reduction in the neutralisation of the omicron strain by corresponding antibodies. Such observations suggest that introducing the point mutations followed by molecular docking may not give the correct picture as reported previously^27^.

**Figure 1.**
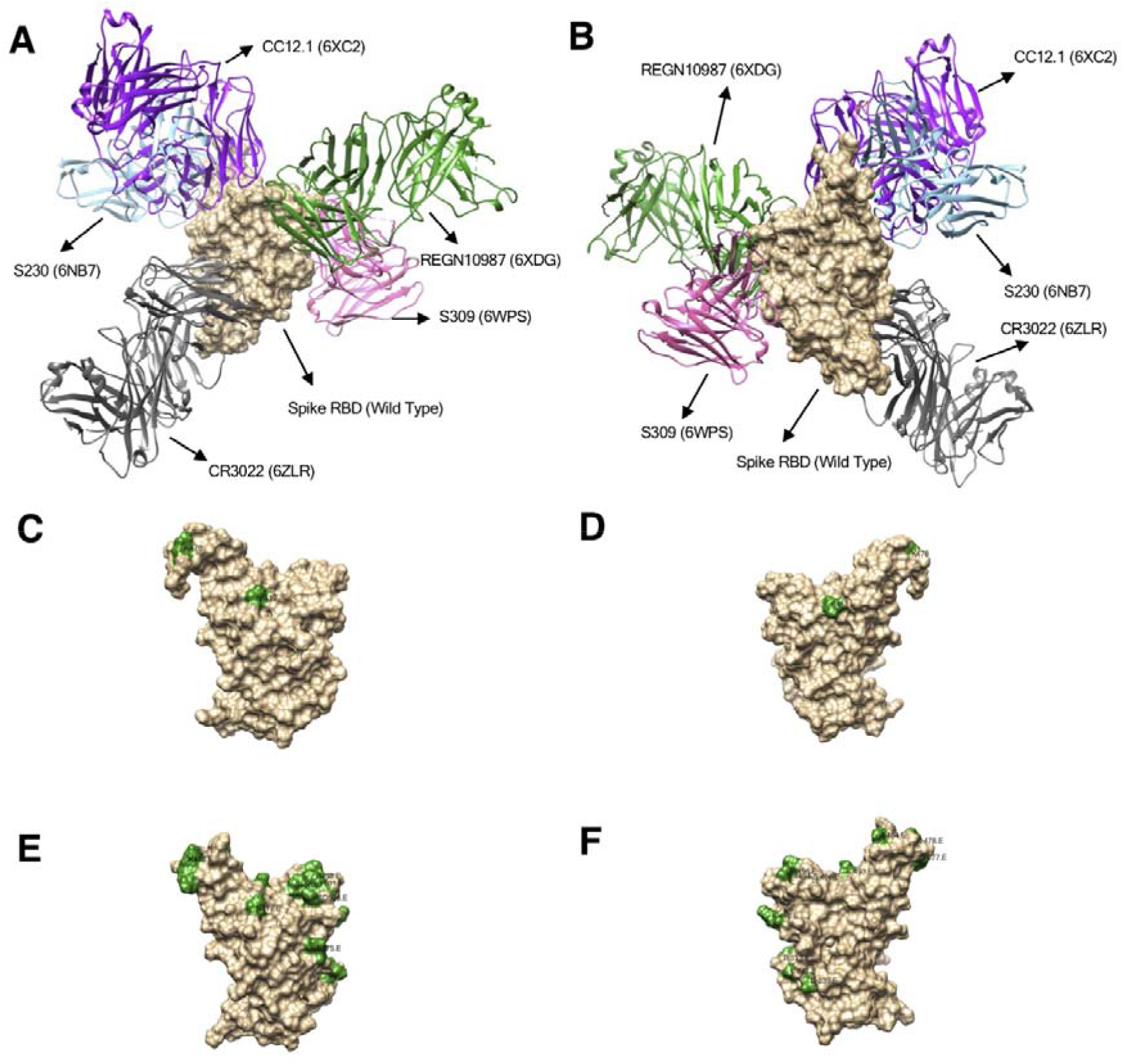
Surface plots showing crystal structure of the spike protein RBD (wild type) in complex with five antibodies binding at their respective binding sites (epitopes) (A) Front view and (B) Rear view for the wild type strain. Surface plot showing mutated residues highlighted in green colour, for Delta (C. Front and D. Rear view) and Omicron (E. Front and F. Rear view). Scientific names of the antibodies along with PDB IDs are /annotated.

Due to the notable mutations acquired by the viral structure, considerable structural changes were expected in the receptor binding epitope of the spike protein. To account for the structural changes firstly point mutations were introduced into the spike RBD structure, as per those present in the delta and omicron variants, and their energies minimised. We observed that static structures do not provide the full picture (Table S1) as the mutations can lead to a partially modified fold and performed molecular dynamic simulation studies on the spike RBD structures, which allowed them to acquire a more reliable adopted dynamic conformation.

After the MD simulations of the spike RBDs, we extracted the most populated structure for all the three strains using the gromos clustering method. Such an approach overcomes the biases towards the end structure and represents the ensemble of most visited conformations during the MD. Superposition of starting and the most populated structure along with the location of the mutations are shown in Figure 2. It is clearly visible that the relaxed (wild strain) and mutated structures (Delta and Omicron strains) exhibit some conformational changes. These changes were more prominent at the epitopes related to the binding of some of the antibodies. Overall wild to delta to omicron variants suggest an increased stability to the viral protein via the attained mutations. We also compared all the three structures after MD (Figure 2D) visualising the differences across them and noticed most visible changes at the mutational site.

**Figure 2.**
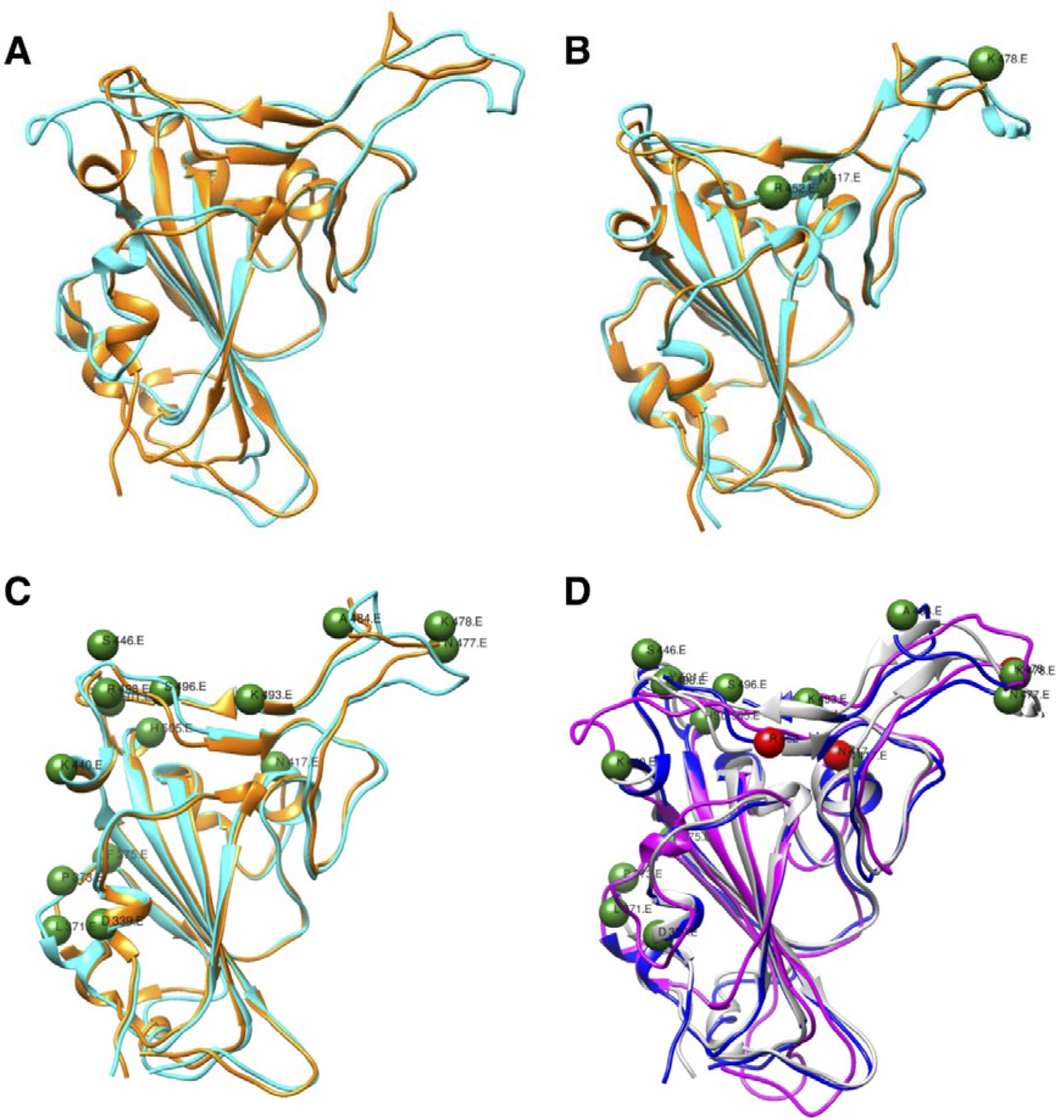
Superposed structures of spike RBD before (orange) and after (cyan) MD simulations in the strains (A) Wild Type. (B) Delta (C) Omicron. Location of mutated residues are depicted by green colour spheres. (D) Superposed structure of the wild type, delta and omicron spike RBD depicting impact of mutations on the protein structure (Magenta = Wild Type, Grey = Delta and Blue = Omicron variant. C-alpha atoms of mutated residues are shown in red (for delta) and green (for omicron) colour spheres.

Represented structures after MD were then subjected to molecular docking analysis. We examined the binding affinity of the protein-neutralising antibody complex due to induced mutations especially after considering the conformational changes on the protein structures during MD simulations. The best conformations generated from the docking analysis obtained by the ClusPro 2.0^33^ were visualised and comparative data was interpreted highlighting the phenomenon of escapism of the mutated SARS COV-2 virus particles from neutralisation by the antibodies due to mutations at the binding sites of the protein.

Comparative docking scores of all the three variants were tabulated in Table 1. The scores depicted a considerable change in the spike RBD-antibody interactions among the variants as compared to the wild type. Especially the omicron variant was showing a significant reduction in the ClusPro 2.0 score as compared to the delta variant. It indicated the impact of the location of mutating residues in the spike RBD. The delta variant did not exhibit considerable difference compared to the wild strain. Two of the antibodies S230 and CC12.1 had slightly compromised neutralisation efficiency while S309 and REGN10987 displayed marginally better binding with the spike RBD compared with the wild strain. ClusPro 2.0 score of CR3022 antibody practically remained unchanged. It is important to mention that despite the constrained antibody-antigen docking, the S230 antibody found other binding epitopes compared with the crystal structure as displayed in Figure S1B. Such changes can be attributed to the observed mutations in Delta strain and relative weak binding of S230 compared with the other antibodies. Overall our results clearly explained why no significant changes in the neutralisation efficiency of the antibodies were observed for the delta variant which is in line with available literature^34^.

**Table 1:** ClusPro 2.0 spike protein-antibody docking scores for the various antibodies.

| **Sr. No.** | **Antibodies (PDB ID)** | **Wild** | **Delta** | **Omicron** |
| --- | --- | --- | --- | --- |
| 1 | S230 (6NB7) | -316.1 | -284.2 | -284.2 |
| 2 | S309 (6WPS) | -754.1 | -846.4 | -506.1 |
| 3 | CC12.1 (6XC2) | -1023.4 | -967.3 | -599.6 |
| 4 | REGN10987 (6XDG) | -856.6 | -977.1 | -570.1 |
| 5 | CR3022 (6ZLR) | -1036.4 | -1034.6 | -606.4 |

The Omicron variant displayed totally different pictures after performing the docking study with the representative structure (most populated during MD). For all the antibodies considered in this study, a notable difference was observed when compared with the wild type (Table 1). S230, S309, CC12.1, REGN10987 and CR3022 display 10, 33, 41, 35 and 41 percentage reductions in their neutralisation capability respectively. Similar reduction was reported in many experimental reports^36^ but as per best of our knowledge none of them explained the molecular mechanism in atomistic level which is discussed below. It is important to highlight that for all the antibodies either other epitopes and/or a different binding pose was observed compared with the crystal structures (Figure S1). The S230, S309 and CR3022 antibodies were binding at different locations while the remaining two displayed different binding poses. Such significant changes for the antibodies in terms of docking scores and binding locations correlated very well with their epitopes and positions of mutated residues in Omicron strain.

CC12.1 exhibits more than 40% reduction in the docking score mainly due to major mutations happening near the receptor binding epitope of spike RBD, thus, aiding mutated variants to escape the neutralising antibodies.

### Characterisation of the antibody-spike RBD binding interface

A closer look at the spike RBD/CC12.1 interface (Figure 3) illustrates that many interactions which were present in the wild type vanished and were replaced by new interactions formed between the spike RBD and the antibody, due to the impact of mutations near the receptor binding site which itself is also the epitope for CC12.1. Interactions formed between the spike RBD residues R403, L455, F456, Y473, A475, E476, F486, N487, Y489 and K493 with the antibody CC12.1, have only been retained in the delta variant compared to the wild type, from which R403 and F456 are located in the vicinity of the mutated residues (Figure 3 & 4).

**Figure 3.**
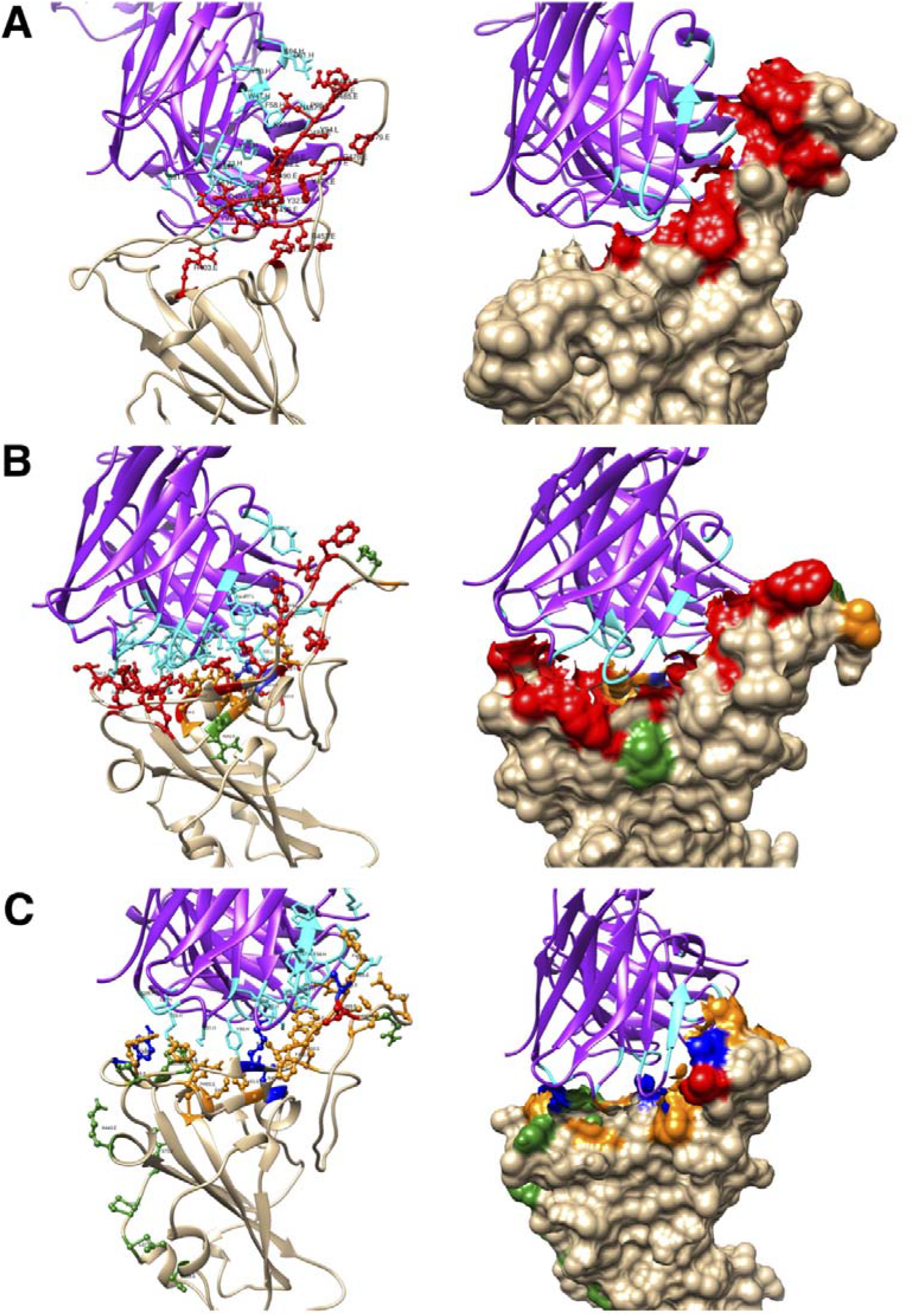
Ribbon and surface diagrams showing the interface region of interaction between the spike RBD (tan) and the neutralising antibody with its heavy and light chain (violet) complex (for antibody CC12.1) for (A) Wild Type (B) Delta and (C) Omicron variant. Interacting residues of the spike RBD, mutated residues and the antibody interacting residues are displayed in red, green and cyan colour respectively. RBD residues which are mutated and interact with antibodies are shown in blue. Interacting residues present in the vicinity (6 Å) of the mutated residues of the spike RBD are shown in orange.

**Figure 4.**
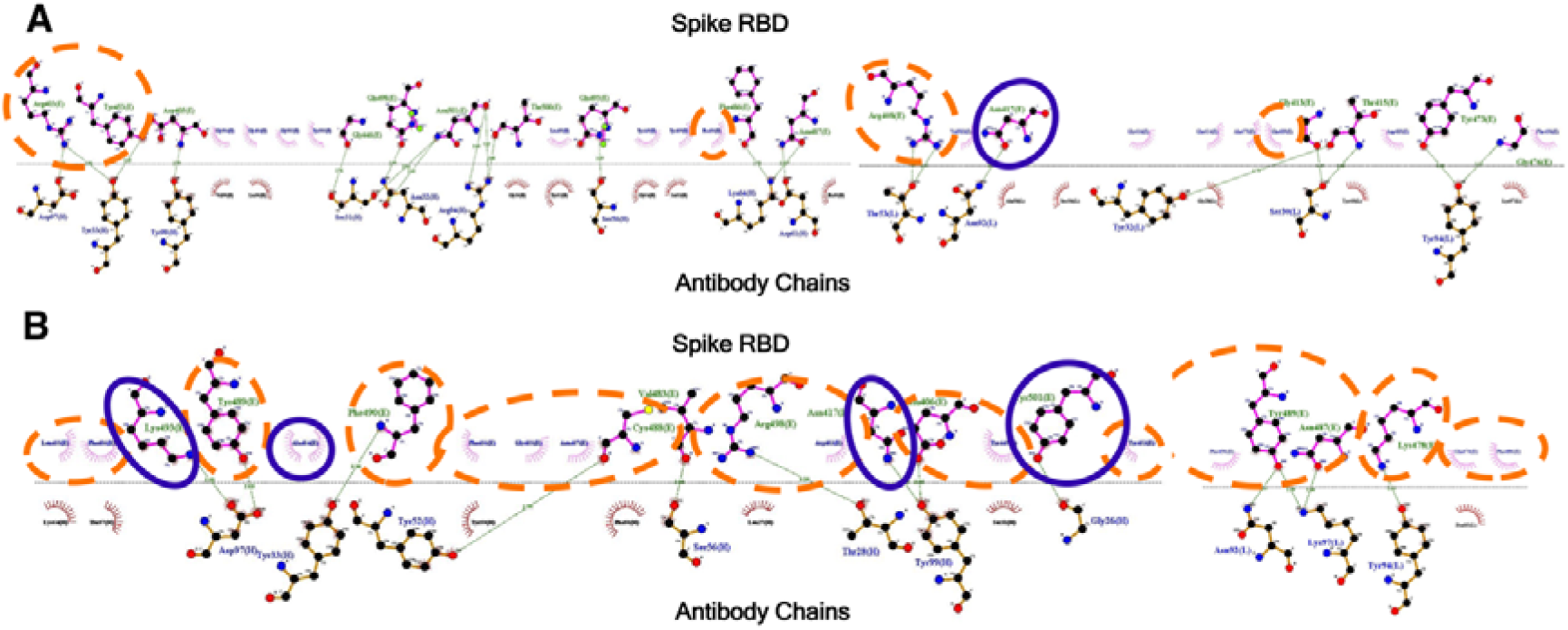
2D plot showing the interface region of interaction between the spike RBD and the neutralising antibody CC12.1 for (A) Delta and (B) Omicron variant. RBD residues which are mutated and interact with antibodies are highlighted by a blue circle. Interacting residues present in the vicinity (6 Å) of the mutated residues of the spike RBD are encircled in orange.

Drastic impacts of mutations were clearly visible in the Omicron variant (Figure 3C). The probable reason points out to be related to the mutations induced in the vicinity of the receptor binding epitope region of the spike RBD, which probably have led to vanishing of some of the previous interactions formed between the spike RBD and the antibody chains (Figure 3 & 4B) and were partially replaced by some new interactions forming but not being able to compensate for the old ones resulting in a notable decrease in the binding score of the omicron in comparison to the wild type. On the contrary, only few of the interacting residues of the spike RBD had an impact on the mutations for the delta strain as majority of them were not in the close proximity of mutated residues (Figure 3B and 4A). Thus, no significant change in the docking score of the delta variant compared to the wild type (Table 1) could be noticed. It was observed that for the omicron strain, spike RBD residues that participate in interactions with neutralising antibodies were either mutated residues (blue circle) or in the close proximity with the mutated residues (orange circle) thus having a significant influence of mutations (Figure 4B). Such observation was majorly absent for the delta strain. Thus, suggested a probable easy escape of the mutated omicron spike protein from coming in recognition by the antibodies and forming a neutralisation complex. A similar trend was observed in the case of the antibodies S309, REGN10987, CR3022 and to some extent for S230 as shown in (Fig S2 to S8). Overall a thorough structural analysis explains the effect of mutations on the antibody-antigen interaction interface and is very well correlated with the docking data.

### Molecular Dynamic (MD) Simulation analysis to study the dynamic nature of the Antibody-Spike RBD Interface

Our findings motivated us to analyse the representative complexes via more sophisticated techniques. Molecular docking largely neglects both the protein flexibility and solvent related terms which are crucial in our case, so we decided to carry out Molecular Dynamics (MD) simulations, as they provide flexibility to proteins besides mimicking the cell-like environment. As MD is computationally expensive we have picked one representative RBD-Antibody complex and performed the comparative MD for Wild Type (WT), Delta and Omicron. The 6XC2 complex was chosen based on substantial change in docking score and its binding interface that involved many mutated residues.

To measure the deviation between positions of an atom with respect to the starting structure, Root Mean Square Deviation (RMSD) values of all simulated structures were evaluated for 300 ns simulation time. RMSD profiles of C-Alpha atoms of the spike RBD for all the three systems (n=3) are displayed in Figure 4A. It was observed that the RBD of WT strain exhibits highest flexibility (~0.55 nm) followed by Delta (~0.4 nm) and Omicron, which shows relatively rigid structure (~0.2 nm) in all three sets of simulations. Higher flexibility of the WT strain allowed it to bind with different antibodies by adjusting its three dimensional structure and maintained that the neutralisation is still feasible by the antibodies. Similarly, the Delta variant also exhibited flexibility to some extent that allowed it to be neutralised by many antibodies however in some cases its affinity decreased as observed in the molecular docking results. Importantly, the Omicron variant displayed relatively rigid structures that must be the after-effect of the large number of mutations. As antigen-antibody interaction is governed by a delicate balance of rigidity and flexibility^35^, this was lost in the case of the Omicron variant leading to a significant reduction of the binding score for the omicron variant (Table 1).

To characterise the interaction interface of the spike RBD and the antibody, Solvent Accessible Surface Area (SASA) of the RBD/antibody interface was calculated. Averaged distributions from all the simulations are displayed in Figure 5B and 5C. It is clearly evident that WT and Delta strain had higher SASA available from both interface sides compared to the Omicron strain. It indeed confirms that both strains can interact with the antibodies with higher affinity further helping in neutralisation due to a large available interface to form antigen-antibody complexes.

**Figure 5:**
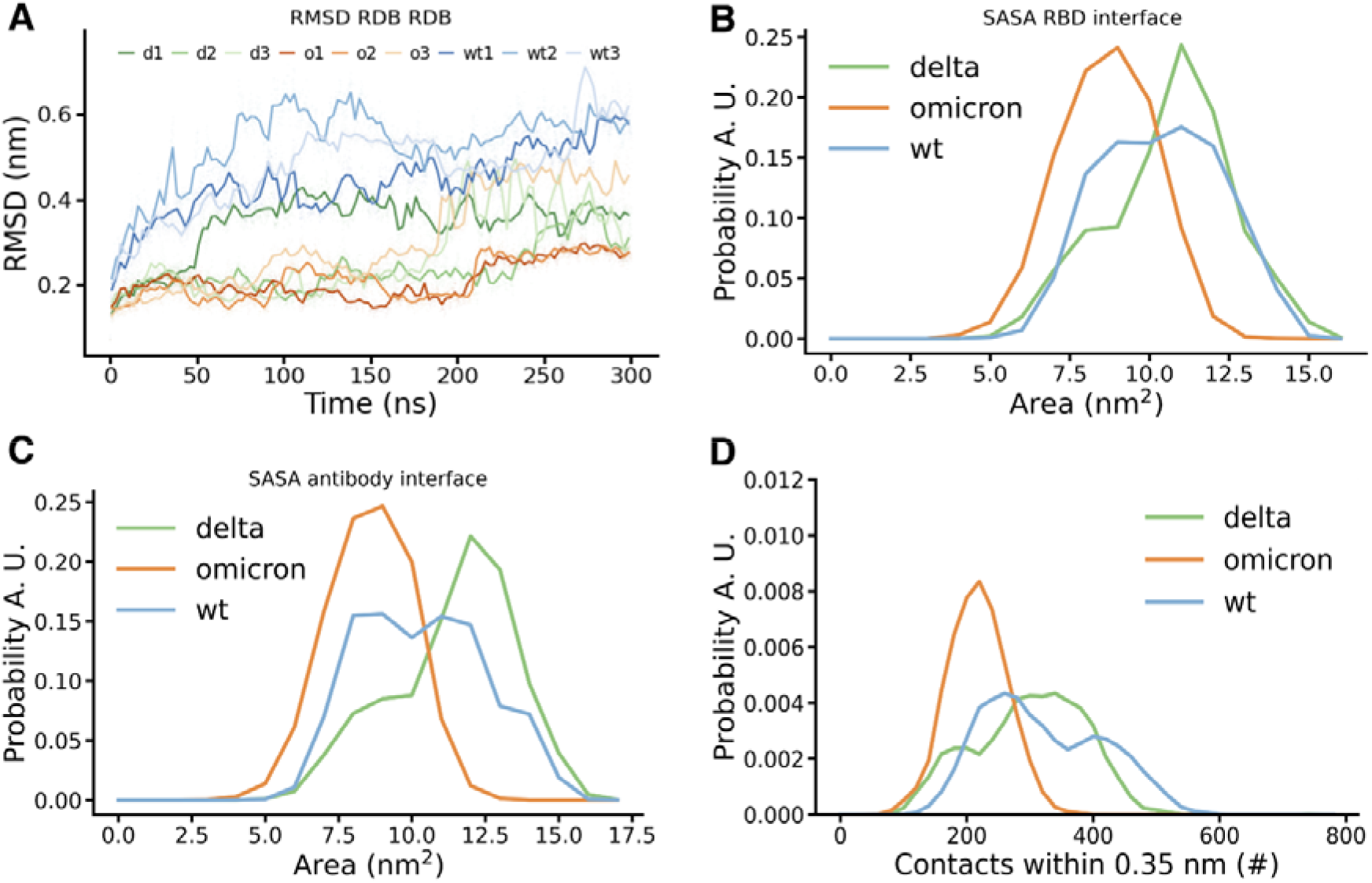
(A) RMSD profile, distributions of (B) SASA from RBD interface, (C) SASA from antibody interface and (D) number of contacts. WT, Delta and Omicron strains are displayed in blue, green and orange colour respectively.

Additional quantification of the interaction strength is provided by the number of contacts between the spike RBD and the antibody. As expected both, WT and Delta strain, form almost similar numbers of contacts with marginally edge to WT strain (Figure 5D). On the other hand, the Omicron strain displayed a ~50% reduction in total number of contacts, which correlated well with the docking score. It clearly depicts the effect of the large number of mutations for the Omicron variant. In the next step, we classified the residues involved in these interactions from the RBD and the antibody perspective. Residues which form contacts for at least 50% of the simulation time during the last 75% of simulations (100 to 300 ns) are illustrated in Figure 6A and 6B. Depicted are residues involved from the RBD and antibody interface along with their probability of being in contact with the counterpart protein respectively.

**Figure 6:**
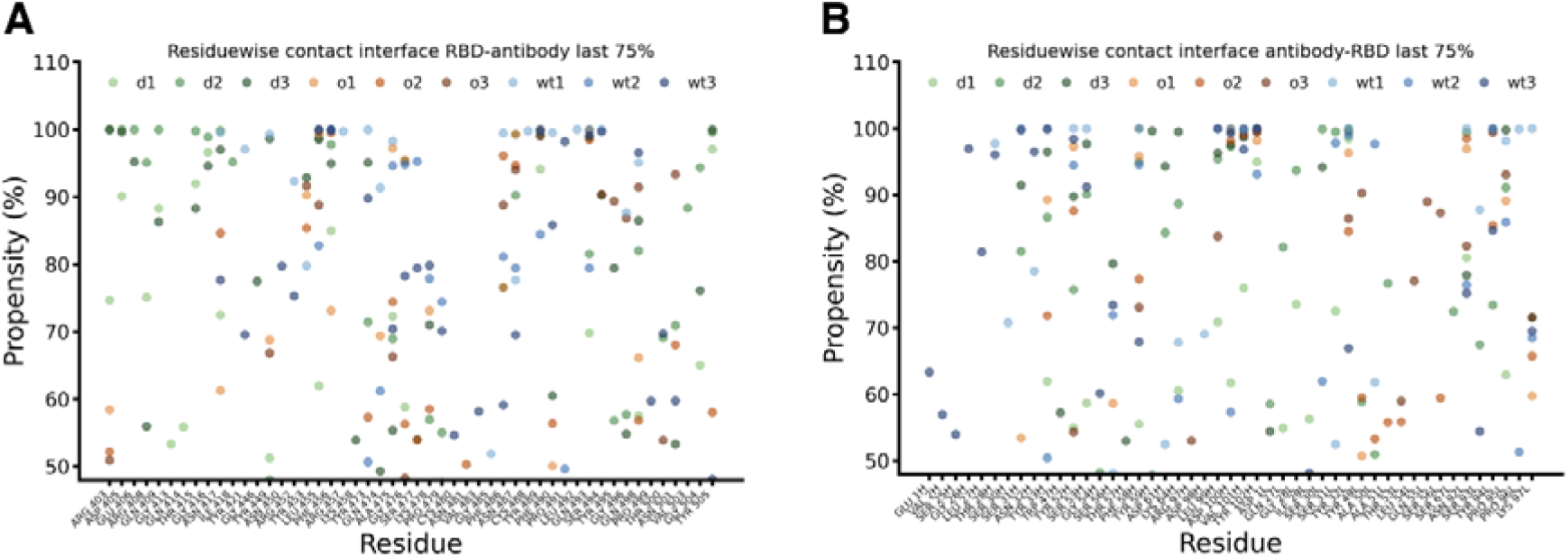
Residue wise contacts and their stabilities during last 75% simulation time for (A) RBD interface and (B) antibody interface. WT, Delta and Omicron strains are displayed in blue, green and orange colour respectively. Data for all three sets of simulations are displayed separately.

With no surprise, both WT and Delta strains were majorly involved in very stable contacts as largely blue and green coloured dots were observed in the range of 80-100% stable contacts. It is important to mention that residues involved in such contacts were distinct in WT and Delta strain. It suggests that due to mutations in the spike RBD the antibody binds at the Delta variant in slightly different locations at RBD compared to the WT but with a similar number of stable contacts. This observation explains that despite the observed mutations in the Delta strain the overall binding is not affected due to lost interactions being compensated by new ones. Therefore, antibodies are able to neutralise the Delta strain. The effect of mutations in the Omicron strain was clearly visible, as except for a few isolated cases, most of the identified residues formed only for 50-70% of the time stable contacts. Importantly, any distinct pattern for the Omicron variant like for Delta and WT was not observed. Here the interacting residues were distributed throughout the interface with reduced stability. Such observations provide the atomic level explanation why the Omicron variant is not able to be neutralised by the majority of antibodies as reported in the literature^24^. A detailed description of the interacting residuals along with their stability is tabulated in Tables S1A/B. Notable interactions which were lost or form less stable contacts in Omicron with respect to WT were observed for residues N417, Y421, R452, R457, K458, Y473, Q474, S477, P479, F490, S494, Q498. It is important to mention that many of these residues are mutated residues or in their close proximity. Similarly we also characterised interactions that are lost or become less stable in Delta compared with WT. However, as discussed above delta formed some new interactions as summarised in the Tables S2A/B that were able to compensate for the lost ones. Other studies have also reported some of these interactions based on cryo-EM structures and neutralization assays^36,37^. However, as per best of our knowledge none of the studies reported all the interacting residues in a dynamic environment and characterized the interactions which are specific to WT, Delta and Omicron.

### End state free energy evaluation via MM-PBSA/GBSA methods

Finally we have quantified the binding energy of antibody-antigen complexes using two approaches. In the first approach we have calculated the average MM-PBSA free energy throughout the trajectory by considering structures at every 10 ns (Table 2). In the other approach we have identified the most populated structure during the MD simulation using the gromos clustering method and calculated the free energy for all three complexes averaged over all three sets of simulations (Table S3).

**Table 2:** Average end state free energies (MM-PBSA) for all the variants during three independent simulation runs.

| Simulation run | WT<sup>1</sup> | Delta<sup>1</sup> | Omicron<sup>1</sup> |
| --- | --- | --- | --- |
| I | -58.31 ± 9.78 | -37.82 ± 12.67 | -36.14 ± 8.75 |
| II | -36.26 ± 7.94 | -49.36 ± 12.69 | -22.57 ± 10.85 |
| III | -39.64 ± 11.81 | -53.99 ± 15.07 | -28.35 ± 8.88 |
| Average | -44.74 ± 9.85 | -47.06 ± 13.48 | -29.02 ± 9.5 |

<sup>1</sup>Energy values are reported in kcal/mol along with the standard deviation.

End point free energy value reiterates the fact that WT and Delta variants have similar binding affinities as observed in molecular docking, number of contacts and SASA. Contrary, the Omicron variant displays a significant reduction in binding energy by 35 to 40%.

The in-depth analysis of the mutations at the interfacial residues and its effect on the binding with the neutralising antibodies across the major dominant variants helps in the designing of the consensus based immunogens where the highly mutable and critical residues could be excluded in the peptide based immunogen sequence. Such immunogens are expected to elicit broadly neutralizing antibodies which may work against the future variants as well^38^. Similarly, the analysis of antibody–antigen contact surfaces using computational tools could be used to guide for the choice of mutations for modelling the antigen-antibody complexes and the rational affinity engineering of therapeutic antibodies^39^.

## Conclusions

The systematic *in silico* docking and MD simulation analysis reveal the impact of the mutations at/near the antibody-antigen interfacial complex. They induce a diminishing affinity of the antibodies towards the omicron mutant by a factor of 35-40%, with an observed reduction, in the total solvent accessible spike RBD surface area followed by an equivalent decrease in the docking score, where the majority of the viral RBD residues at the interface have mutated. This severe change in the binding epitope aids the viral spike protein of the omicron variant from escaping the antibody neutralisation. For the delta variant, with only limited mutations near/at the viral interface, only a weak effect of the mutations has been observed, causing only locally limited distortions in the binding pattern which are easily compensated by the neighbouring residues. This analysis shines light on important aspects necessary for the development of a robust and effective vaccine and immunisation in its truest essence. The applied *in silico* methods can be a fast and economical strategy for the early prediction of the effects of newly emerging SARS-CoV-2 variants at the molecular level and provide first clues to the researchers for further investigations.

## Material and methods

### Protein selection and structure preparation

#### Spike Protein

The receptor binding domain (RBD) of the spike protein (S1) was selected for the study, as the major population of the serum neutralising antibodies target the RBD domain of the spike protein, supported by various clinical and serological studies done on the COVID-19 infected patients^40^.

The three dimensional structure of the wild strain was retrieved from Protein Data Bank^41^ (http://www.rcsb.org) with PDB ID: 6M0J^42^. The co-crystallized protein ACE2 in the model was separated through UCSF Chimera^43^, along with the other non-interacting ions and water molecules before performing the molecular docking.

#### Antibody Structures

The epitope sites on the RBD of the spike protein suggested the presence of three binding sites where the antibodies having neutralisation ability bind to, namely^44^: (i) receptor binding site; (ii) CR3022 cryptic site; and (iii) S309 proteoglycan site. Based on this knowledge, the neutralising antibodies which bind to the distinct epitope sites of the spike RBD were selected - the S230^29^ (PDB ID: 6NB7) and the CC12.1^30^ (PDB ID: 6XC2) binding the receptor binding site region, the REGN10987^31^ (PDB ID: 6XDG) and S309^32^ (PDB ID: 6WPS) attaching at the proteoglycan site^44^ and the ones binding the CR3022^28^ site (PDB ID: 6ZLR).

#### Inducing mutations

The major notorious clades of the mutated strains are - the Delta (B.1.617.2) and the Omicron (B.1.1.529) variants, which has led to the major havoc in the world recently.

For creating the mutated clades of the other variants of SARS COV-2, the mutations were introduced at the specific residual locations in the RBD domain. K417N, L452R and T478K in the RBD of the delta variant and G339D, S371L, S373P, S375F, K417N, N440K, G446S, S477N, T478K, E484A, Q493K, G496S, Q498R, N501Y and Y505H in that of the Omicron variant. The point mutations were introduced with the rotamer feature of UCSF Chimera^43^ by selecting favourable rotamers. Subsequently the resulting structure was energy minimised to reduce/minimise still possible clashes/contacts of the chains in the mutated residues with that of the surrounding residues in the protein.

### Molecular docking

The study was conducted using ClusPro 2.0^33^, by docking the spike RBD protein of the Wild strain and the mutated strains B.1.617.2 (Delta) and B.1.1.529 (Omicron) with the various antibody proteins.

For protocol validation, the crystalline protein structure of SARS-CoV-2 spike RBD in complex with neutralising antibody CC12.1 was selected. The two proteins were detached from each other and redocked at their interacting site. The changes, in the observed binding pattern, were negligible, and the estimated root mean square deviation (RMSD) value of 0.832 Å, was in a clearly acceptable range, thus, validating the docking protocol.

The interaction residues and binding sides were selected based on the literature^28–32^, together with visualisation and identification of the molecular interactions among the spike RBD and the antibodies using UCSF Chimera and LigPlot+^45^. During the docking process, antibody mode was selected for generating the best clusters after docking analysis and complementarity-determining regions (CDRs) were masked for effective antigen-antibody complex formation^46^. Best models were selected according to the highest Cluster member size generated and therefore having the optimum score. The collective scores of all the docked complexes were compared and the results were interpreted.

Since the static structures provide only superficial information as the mutations can lead to a partially modified fold, the wild and the mutant spike RBD structures were subjected to molecular dynamic simulations studies to provide more reliable adopted dynamic conformations. All the strains were subjected to 100 ns of MD simulations (as discussed below) and representative structures were extracted for further docking studies. These representative structures were obtained by clustering of the MD simulations with the gromos algorithm together with a cut off distance of 0.2 nm and correspond to the cluster centroid of the biggest obtained cluster.

Complex structures after the molecular docking were visualised using UCSF-Chimera^43^ and 2D plots were generated using Lig-Plot +^45^.

### Molecular dynamics (MD) Simulation studies

All molecular dynamics simulations were performed with GROMACS version 2021.4^47^. We used the CHARMM36m^48,49^; forcefield for the antibody protein CC12.1 (PDB ID: 6XC2)^50,51^ together with the TIP3P water model^52^. The force field parameters for the system have been generated with the Input generator tools in CHARMM-GUI ^51,53,54^ using Solution Builder. The two missing fragments (three and four residues) in the chain H of the antibody part of the structure were built with modeller using the loop modeller routine^55,56^. The initial structures for the Delta and Omicron mutant were obtained by docking the antibody into a representative structure of the RBD, obtained from a MD simulation, with ClusPro 2.0 as discussed above.

The simulation box was set to dodecahedron and defined in such a way that the minimum distance of the structure and the box was at least 1.5 nm for the initial RBD simulation and 2.0 nm for the docked complexes and subsequently solvated with water and neutralised with potassium chloride together with an additional concentration of 150 mmol/L.

The following settings have been applied. The Leapfrog integrator was utilised together with all bonds being constrained by the LINCS algorithm^57^ in order to enable a time-step of 2 fs. We used a modified cut off for short-ranged electrostatic and Lenard Jones interactions of 1.2 nm and applied a switching function to smoothly approach the cut off between 1.0 and 1.2 nm.

Particle mesh Ewald (PME)^58^ method was applied to calculate Long-range Coulomb interactions. The neighbour list was updated every 10 steps. In a first step, all systems were conducted to energy minimisation with the steepest-descent algorithm for 50000 steps. Subsequently two consecutive equilibration simulations followed (100 ps each) in a canonical (NVT) and later on isobaric-isothermal (NPT) ensemble with position restraints on the heavy atoms of the proteins. The final production runs without position restraints were 100 ns long for the initial RBD simulation and 300 ns long for the antibody complexes. The temperature was maintained at 303.15 K with the Nose-Hoover^59^ thermostat applying a coupling time of 1 ps. The bulk systems were simulated in an isobaric-isothermal ensemble, where the pressure was set to 1 atm using isotropic Parrinello-Rahman pressure coupling^60^ with a pressure relaxation time of 5 ps for the system. The simulations of the antibody complexes were performed as triplicates with different initial velocities resulting in three independent simulations.

### Analysis of MD Simulations

To describe the interaction strength of the RDB complexes with the antibody we calculated the following features. Number of contacts, interaction area and residue wise contact probability.

#### The number of contacts between RBD and antibodies

The values have been calculated with the gromacs hbond tool where all atoms within 0.35 nm of both groups were considered as being in contact. We report the distribution over all three simulations for every RBD-Antibody complex, e. g. Wild Type, Delta and Omicron variant.

#### Solvent accessible surface (SASA) of the interface between the RBD and antibody and vice versa

The surface was calculated with the gromacs sasa tool by subtracting the SASA of the RBD part of the RBD-Antibody complex from the SASA calculated for the RBD only. For the other side of the interface the SASA part of the Antibody-RBD complex was subtracted from the SASA of the antibody. Also here we report the distribution of the calculated areas for all simulations.

#### Residue wise probability of the interface amino acids in the RBD and antibody to be in contact with the respective counterpart, RBD to antibody and antibody to RBD

Here we utilised the mindist tool of gromacs and reported the residue wise minimum distance of the RBD amino acids to any atom of Antibody and vice versa, the residue wise minimum distance between the amino acids of the antibody to any atom of RBD. In the plot and the corresponding tables, we report all amino acids which have been in contact (minimum distance of 0.35 nm) to the counterpart in the complex for at least 50 percent of the simulation time. The values are reported for all simulations separately in two plots showing the residue wise contacts for the directions RBD to antibody and the antibody to RBD.

#### Free energy calculations

End state free energy calculations were performed using two different tools and approaches. In the first approach, *gmx_mmpbsa* tools, recently developed by Valdés-Tresanco^61^ and co-workers based on AMBER’s MMPBSA.py script, were used. In this approach, End state free energy was calculated throughout the trajectory with a 10ns interval. We report the energies for all three simulations for every RBD-Antibody complex, e. g. Wild Type, Delta and Omicron variant. This approach provides the energy considering all the conformations observed in MD. In the other approach free energies were calculated using the Prime MM-GBSA^62,63^ module of the Schrödinger Suit. Herein, we have considered representative structures for the performed calculations. The representative structures were obtained by clustering of the MD simulations with the gromos algorithm together with an appropriate cut off distance and correspond to the cluster centroid of the biggest obtained cluster. This approach mimics the molecular docking approach as was considering a single, most populated structure for the energy calculation.

## Supporting information

molecular docking data, the antibody-RBD complexes for other antibodies, the details of interacting residues and the results from MM/GBSA calculations

## Acknowledgements

The authors acknowledge support by the High Performance and Cloud Computing Group at the Zentrum für Datenverarbeitung of the University of Tübingen, the state of Baden-Württemberg through bwHPC, and the German Research Foundation (DFG) through grant no. INST 37/935-1 FUGG. AJ acknowledges the Department of Biotechnology, Government of India for the Ramalingaswami Re-entry Fellowship-2019. Authors acknowledge support by BIT-Mesra for necessary software and HPC resources.

## Supporting Information

The supporting information contains the molecular docking data, the antibody-RBD complexes for other antibodies, the details of interacting residues and the results from MM/GBSA calculations.

## Notes

### Competing Interest Statement

The authors have declared no competing interest.

