## Supplementary material for "Structural basis of Omicron immune evasion: A comparative computational study of Spike protein-Antibody interaction": molecular docking data, the antibody-RBD complexes for other antibodies, the details of interacting residues and the results from MM/GBSA calculations

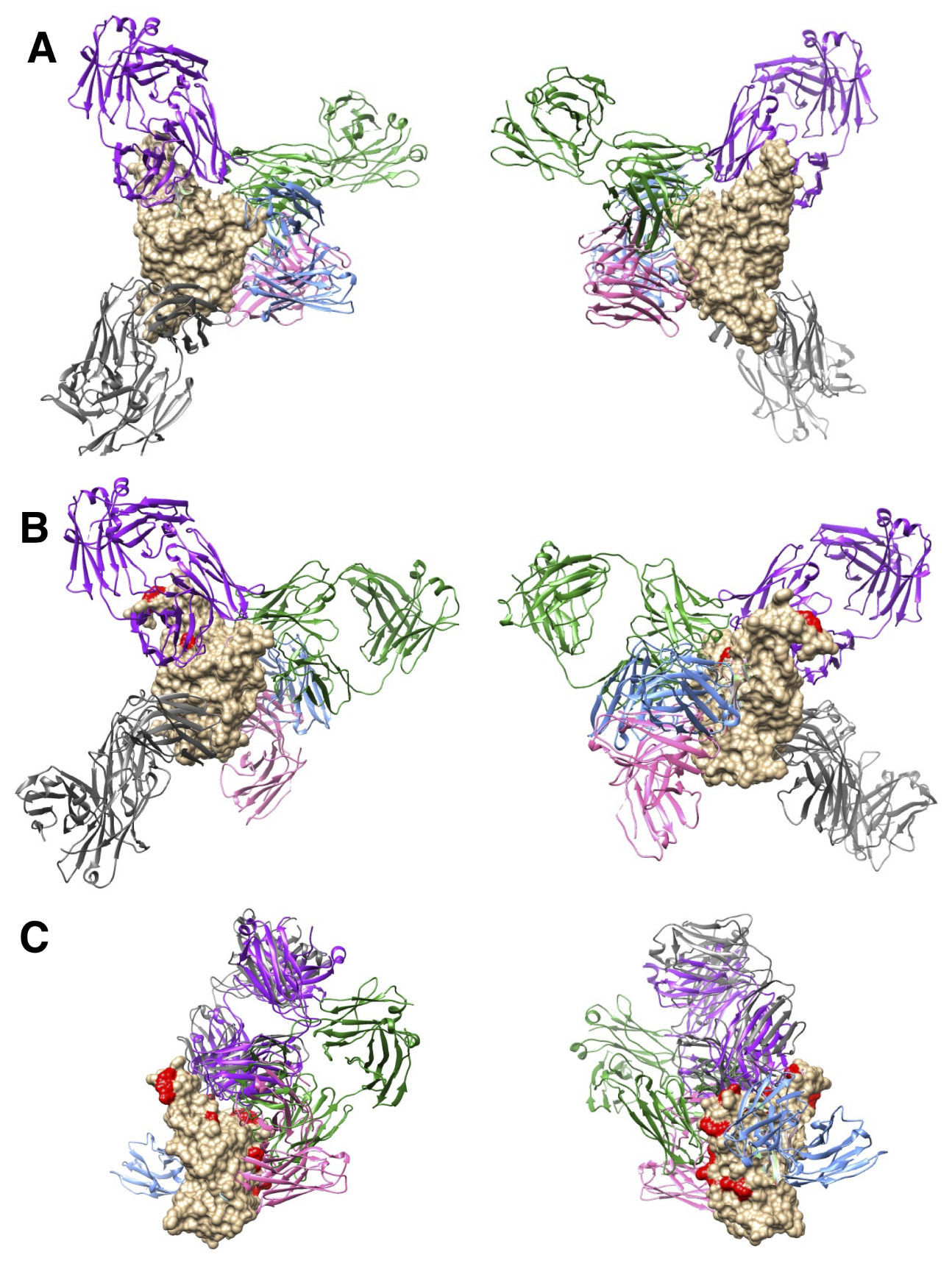

Figure S1. Diagram showing crystal structure of the spike protein RBD. A. Wild type (front and rear) B. Delta (front and rear) C. Omicron (front and rear) in complex with the five antibodies forming complexes after performing the MD simulation and molecular docking analysis (Surface tan plot - Spike RBD, Purple - CC12.1 (PDB ID 6XC2), Blue - S230 (PDB ID 6NB7), Pink - S309 (PDB ID 6WPS), Green - REGN10987 (PDB ID 6XDG) and Grey - CR3022 (PDB ID 6ZLR) Red – Mutated Residues). Scientific name of the antibody along with PDB ID is shown.

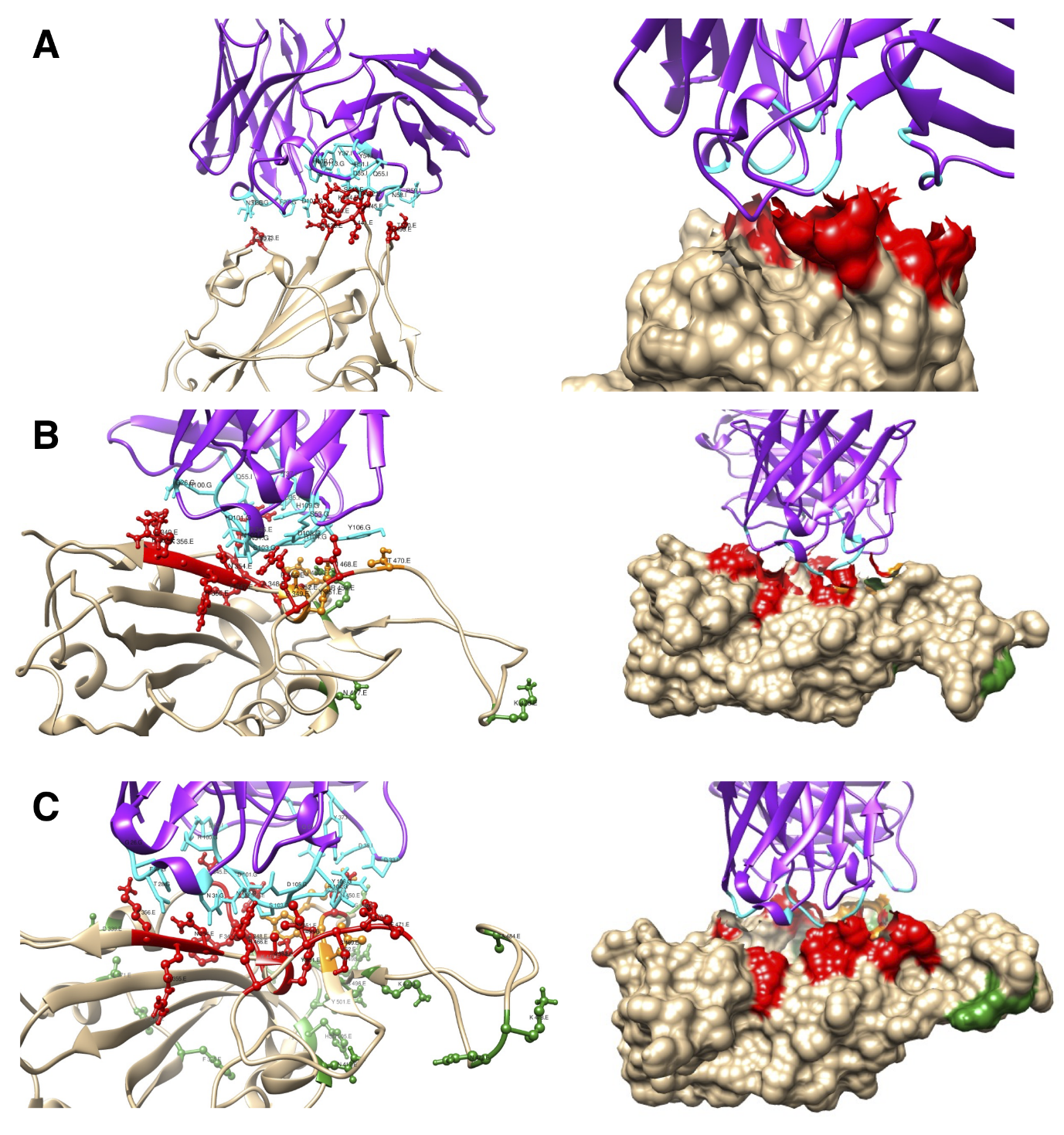

Figure S2. Ribbon and surface diagrams showing the interface region of interaction between the spike RBD (tan) and the neutralising antibody with its heavy and light chain (violet) complex (for antibody S230) for (A) Wild Type, (B) Delta and (C) Omicron variant. Interacting residues of the spike RBD, mutated residues and the antibody interacting residues are displayed in red, green and cyan colour respectively. RBD residues which are mutated and interact with antibodies are shown in blue. Interacting residues present in the vicinity (6 Å) of the mutated residues of the spike RBD are shown in orange.

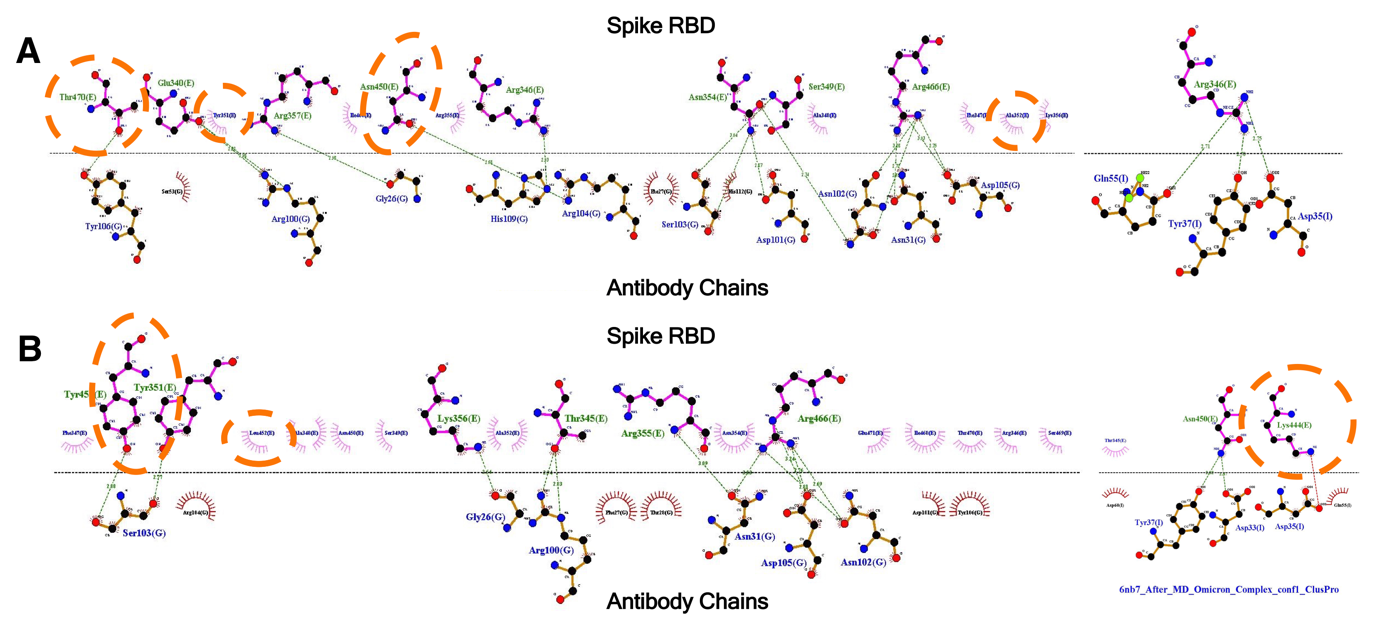

Figure S3. 2D plot showing the interface region of interaction between the spike RBD and the neutralising antibody S230 for (A) Delta and (B) Omicron variant.  RBD residues which are mutated and interact with antibodies are highlighted by a blue circle. Interacting residues present in the vicinity (6 Å) of the mutated residues of the spike RBD are encircled in orange.

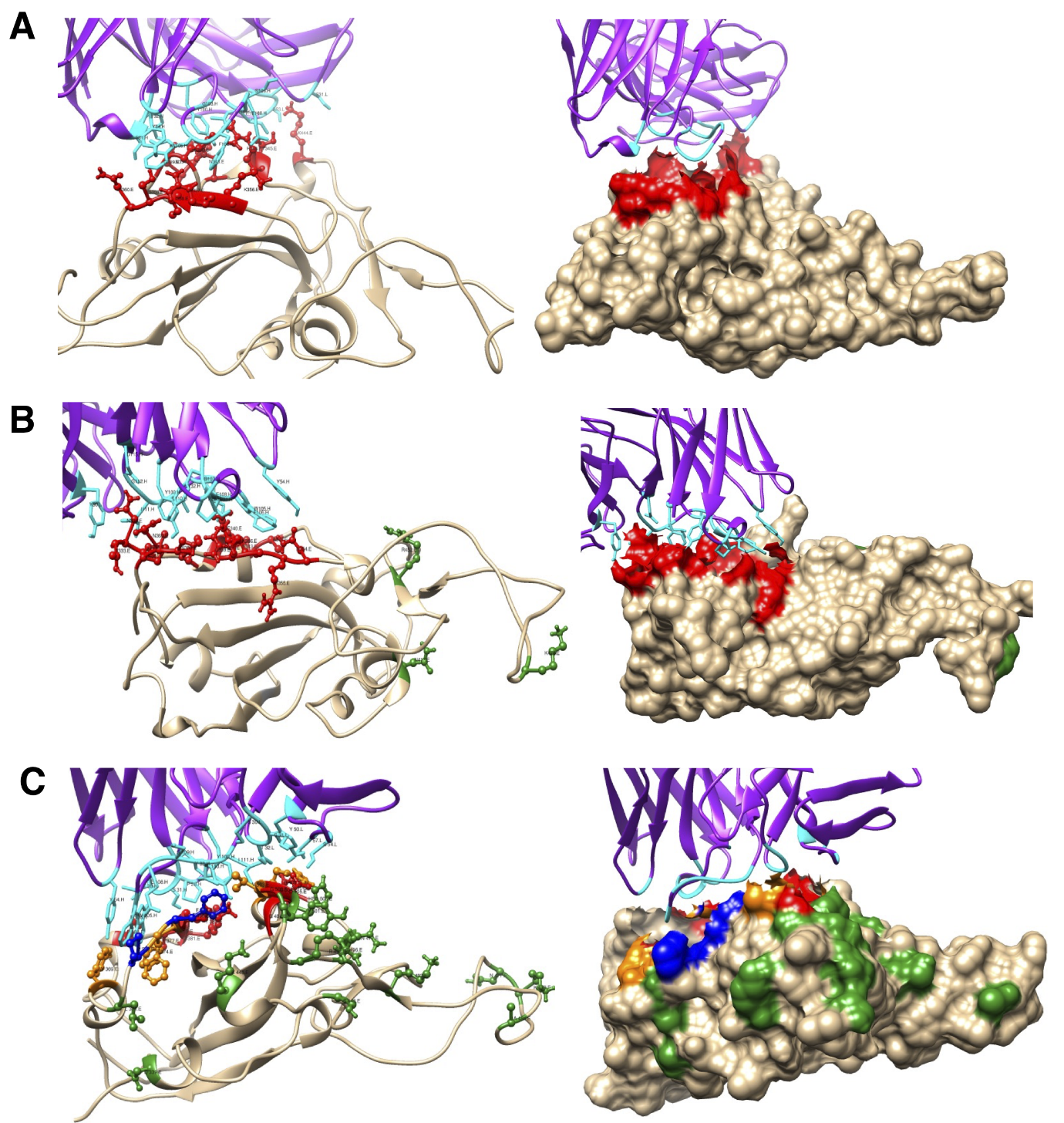

Figure S4. Ribbon and surface diagrams showing the interface region of interaction between the spike RBD (tan) and the neutralising antibody with its heavy and light chain (violet) complex (for antibody S309) for (A) Wild Type, (B) Delta and (C) Omicron variant. Interacting residues of the spike RBD, mutated residues and the antibody interacting residues are displayed in red, green and cyan colour respectively. RBD residues which are mutated and interact with antibodies are shown in blue. Interacting residues present in the vicinity (6 Å) of the mutated residues of the spike RBD are shown in orange.

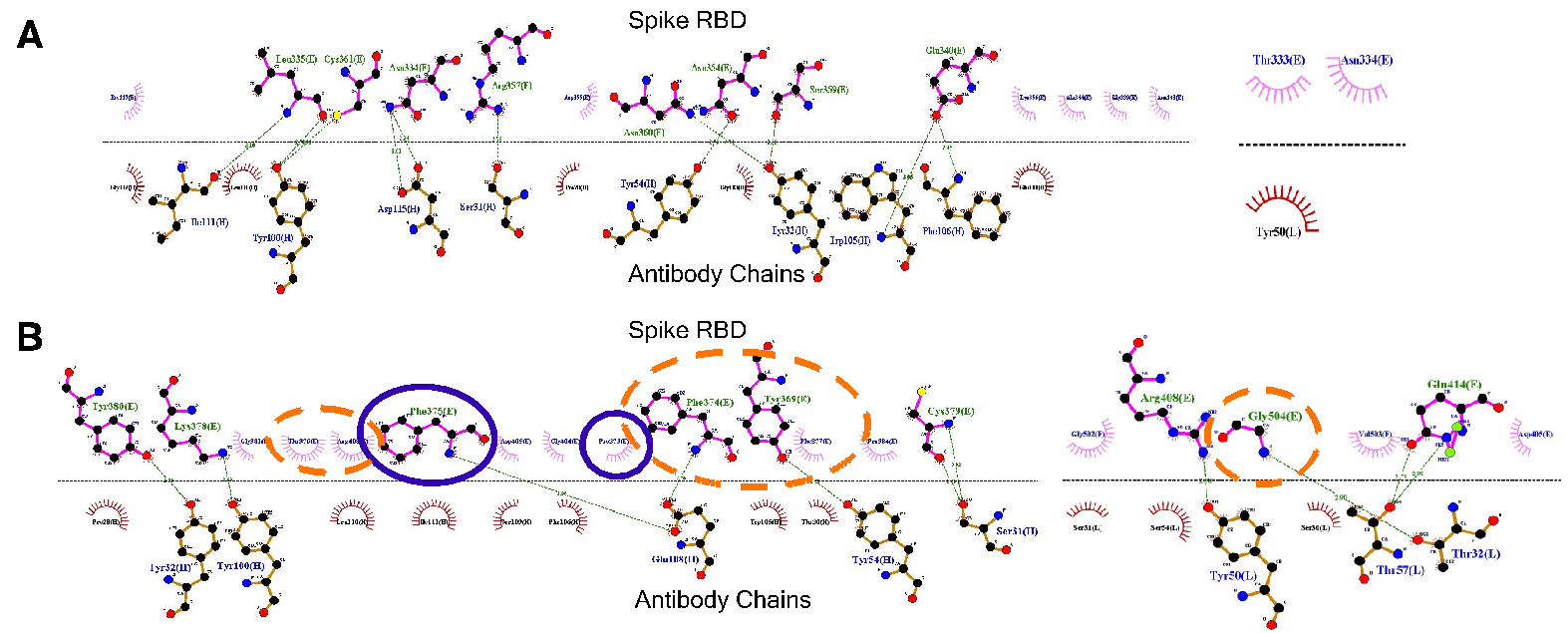

Figure S5. 2D plot showing the interface region of interaction between the spike RBD and the neutralising antibody S309 for (A) Delta and (B) Omicron variant. RBD residues which are mutated and interact with antibodies are highlighted by a blue circle. Interacting residues present in the vicinity (6 Å) of the mutated residues of the spike RBD are encircled in orange.

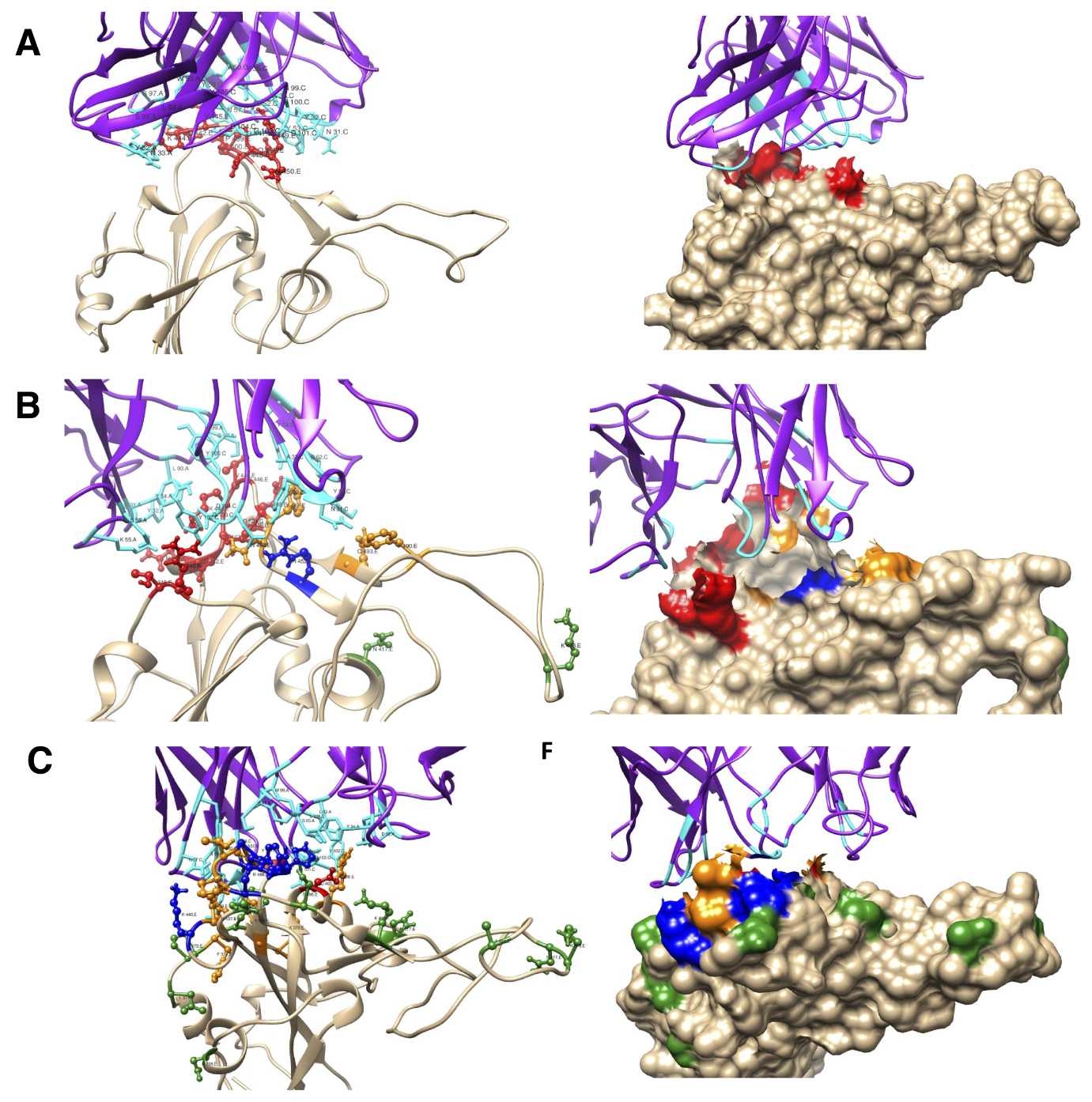

Figure S6. Ribbon and surface diagrams showing the interface region of interaction between the spike RBD (tan) and the neutralising antibody with its heavy and light chain (violet) complex (for antibody REGN10987) for (A) Wild Type, (B) Delta and (C) Omicron variant. Interacting residues of the spike RBD, mutated residues and the antibody interacting residues are displayed in red, green and cyan colour respectively. RBD residues which are mutated and interact with antibodies are shown in blue. Interacting residues present in the vicinity (6 Å) of the mutated residues of the spike RBD are shown in orange.

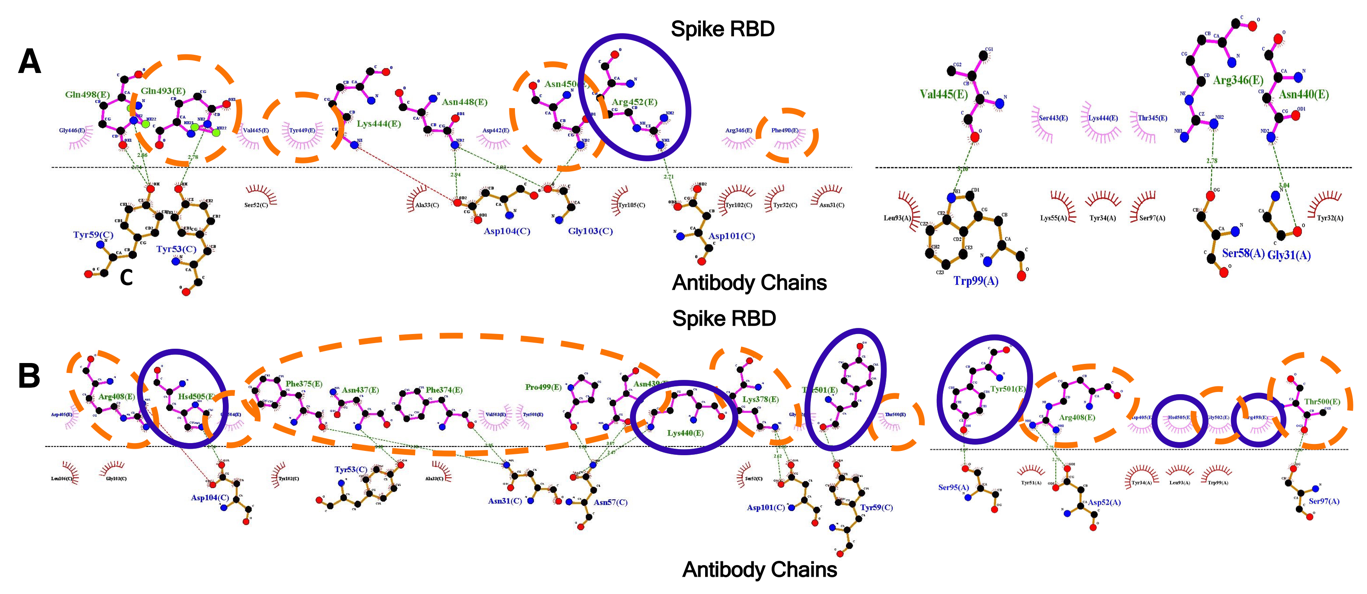

Figure S7. 2D plot showing the interface region of interaction between the spike RBD and the neutralising antibody REGN10987 for (A) Delta and (B) Omicron variant. RBD residues which are mutated and interact with antibodies are highlighted by a blue circle. Interacting residues present in the vicinity (6 Å) of the mutated residues of the spike RBD are encircled in orange.

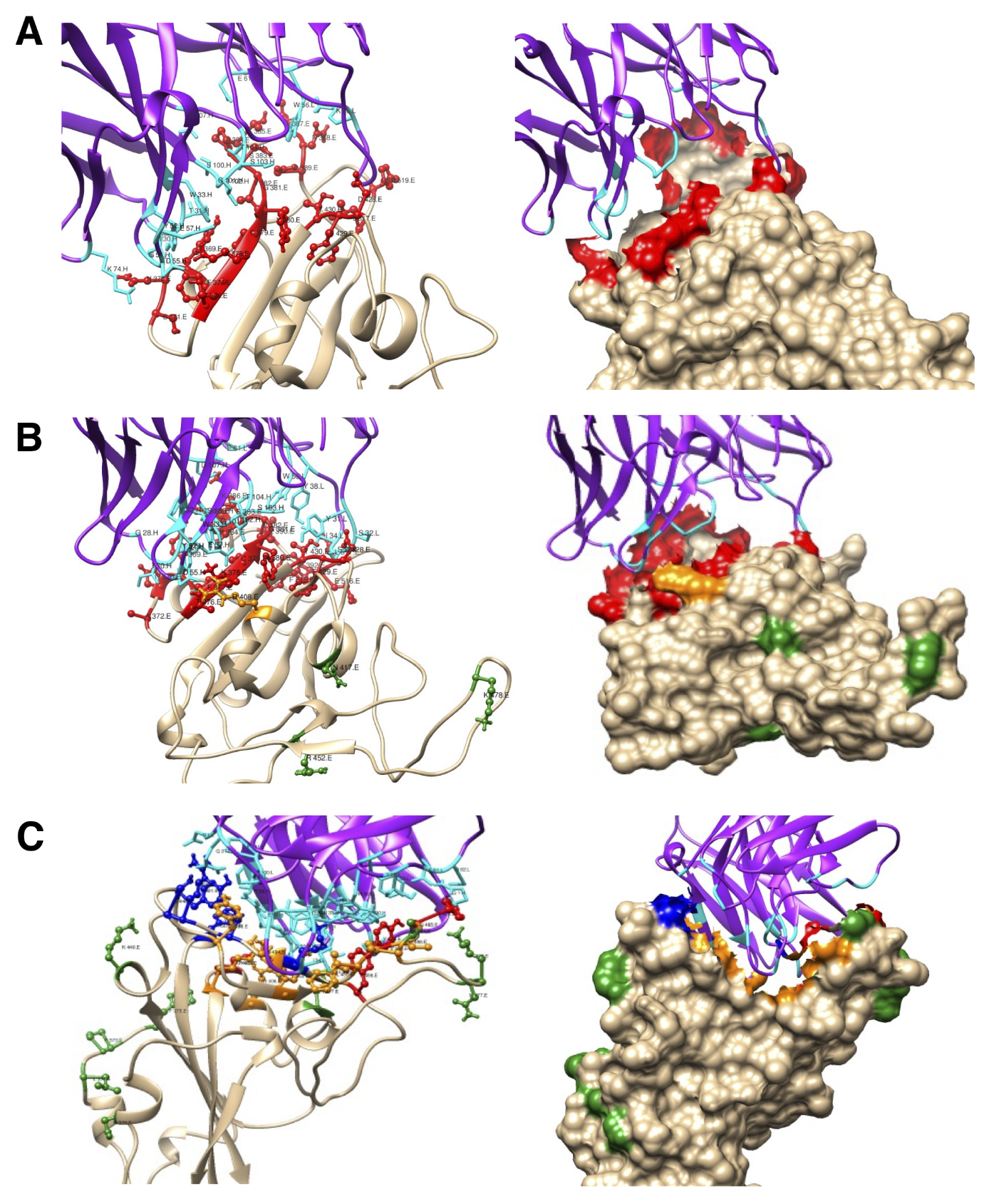

Figure S8. Ribbon and surface diagrams showing the interface region of interaction between the spike RBD (tan) and the neutralising antibody with its heavy and light chain (violet) complex (for antibody CR3022) for (A) Wild Type, (B) Delta and (C) Omicron variant. Interacting residues of the spike RBD, mutated residues and the antibody interacting residues are displayed in red, green and cyan colour respectively. RBD residues which are mutated and interact with antibodies are shown in blue. Interacting residues present in the vicinity (6 Å) of the mutated residues of the spike RBD are shown in orange.

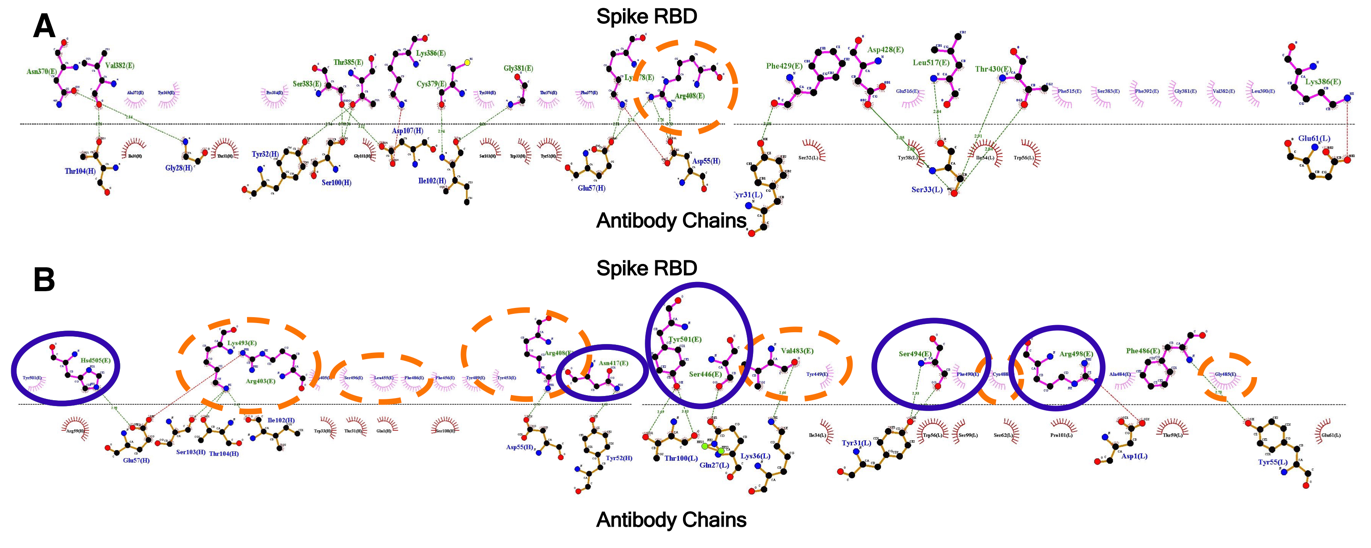

Figure S9. 2D plot showing the interface region of interaction between the spike RBD  and the neutralising antibody CR3022 for (A)  Delta and (B) Omicron variant.  RBD residues which are mutated and interact with antibodies are highlighted by a blue circle. Interacting residues present in the vicinity (6 Å) of the mutated residues of the spike RBD are encircled in orange.

Table S1: ClusPro 2.0 spike protein-antibody docking scores for the various antibodies before MD simulation of spike RBD.

| **Sr. No.** | **Antibodies (PDB ID)** | **Wild** | **Delta** | **Omicron** |
| --- | --- | --- | --- | --- |
| 1 | S230 (6NB7) | -306.6 | -309.5 | -311.5 |
| 2 | S309 (6WPS) | -914.9 | -647.0 | -627.0 |
| 3 | CC12.1 (6XC2) | -1039.8 | -1053.1 | -1052.6 |
| 4 | REGN10987 (6XDG) | -935.0 | -922.3 | -877.5 |
| 5 | CR3022 (6ZLR) | -1127.9 | -1077.3 | -1051.7 |

Table S2A: Residue Wise contacts and their stabilities during the last 75% of the simulation time for the RBD interface.

| **Residue** | **d1^#^** | **d2^#^** | **d3^#^** | **o1^#^** | **o2^#^** | **o3^#^** | **wt1^#^** | **wt2^#^** | **wt3^#^** |
| --- | --- | --- | --- | --- | --- | --- | --- | --- | --- |
| **ARG403** | 74.63 | 100.00 | 99.96 | 58.42 | 52.15 | 50.91 | 7.51 | 1.82 | 0.13 |
| **ASP405** | 90.09 | 100.00 | 99.64 | 8.80 | 18.75 | 2.00 | 0.00 | 0.00 | 0.00 |
| **GLU406** | 35.10 | 99.96 | 95.20 | 22.79 | 10.62 | 0.13 | 1.11 | 0.00 | 0.09 |
| **ARG408** | 75.08 | 95.07 | 55.93 | 9.42 | 3.60 | 0.04 | 0.00 | 0.00 | 0.00 |
| **GLN409** | 88.27 | 99.96 | 86.32 | 0.18 | 0.04 | 0.00 | 0.31 | 0.00 | 0.00 |
| **GLY413** | 53.31 | 0.13 | 0.00 | 0.00 | 0.00 | 0.00 | 0.00 | 0.00 | 0.00 |
| **GLN414** | 55.84 | 18.79 | 0.09 | 0.00 | 0.00 | 0.00 | 0.00 | 1.24 | 0.00 |
| **THR415** | 91.96 | 99.78 | 88.27 | 1.20 | 4.18 | 0.00 | 0.00 | 9.91 | 0.00 |
| **GLY416** | 96.62 | 98.89 | 94.58 | 4.13 | 4.89 | 0.00 | 13.82 | 3.69 | 0.22 |
| **ASN417** | 72.46 | 99.82 | 97.02 | 61.31 | 84.63 | 44.87 | 99.56 | 44.07 | 77.65 |
| **ILE418** | 19.59 | 95.16 | 6.71 | 3.47 | 0.71 | 0.00 | 18.97 | 0.00 | 4.09 |
| **TYR421** | 9.91 | 43.45 | 0.89 | 39.67 | 17.46 | 0.13 | 97.07 | 24.03 | 69.57 |
| **GLY446** | 0.00 | 25.19 | 77.48 | 11.55 | 30.70 | 46.02 | 0.13 | 14.62 | 0.00 |
| **TYR449** | 51.22 | 48.02 | 98.62 | 68.77 | 38.12 | 66.81 | 99.33 | 25.19 | 1.95 |
| **ASN450** | 0.00 | 0.00 | 0.00 | 0.00 | 0.00 | 0.00 | 14.30 | 2.09 | 79.74 |
| **ARG452** | 0.00 | 0.00 | 0.27 | 0.00 | 0.00 | 0.00 | 92.31 | 0.36 | 75.30 |
| **TYR453** | 17.99 | 39.05 | 92.85 | 90.27 | 85.38 | 91.60 | 79.79 | 36.25 | 5.86 |
| **LEU455** | 61.97 | 98.80 | 98.53 | 99.42 | 99.78 | 88.80 | 100.00 | 82.76 | 99.96 |
| **PHE456** | 84.94 | 97.78 | 94.94 | 73.12 | 99.51 | 39.45 | 100.00 | 99.91 | 99.96 |
| **ARG457** | 0.00 | 0.00 | 0.09 | 2.62 | 6.66 | 0.00 | 99.73 | 38.12 | 8.57 |
| **LYS458** | 3.86 | 0.89 | 53.93 | 0.44 | 0.13 | 0.00 | 36.83 | 46.42 | 5.78 |
| **TYR473** | 30.43 | 71.39 | 95.07 | 2.53 | 57.31 | 0.44 | 99.96 | 50.69 | 89.78 |
| **GLN474** | 33.67 | 3.78 | 49.31 | 69.39 | 7.77 | 0.00 | 91.34 | 61.22 | 0.00 |
| **ALA475** | 72.28 | 68.95 | 55.35 | 97.25 | 74.41 | 66.28 | 98.27 | 94.62 | 70.41 |
| **GLY476** | 58.82 | 33.01 | 95.29 | 95.47 | 56.29 | 48.25 | 25.68 | 94.80 | 78.23 |
| **SER477** | 39.18 | 23.68 | 54.02 | 41.45 | 53.93 | 42.83 | 46.11 | 95.25 | 79.43 |
| **LYS478** | 33.81 | 56.95 | 70.99 | 73.12 | 58.51 | 31.36 | 37.94 | 77.88 | 79.83 |
| **PRO479** | 35.76 | 55.04 | 47.67 | 18.04 | 34.21 | 0.76 | 47.13 | 74.41 | 70.10 |
| **CYS480** | 21.77 | 19.15 | 29.81 | 16.53 | 45.58 | 0.62 | 10.31 | 33.10 | 54.64 |
| **ASN481** | 17.64 | 6.66 | 22.39 | 9.82 | 50.33 | 0.27 | 12.57 | 24.21 | 10.40 |
| **VAL483** | 0.04 | 0.04 | 1.07 | 5.38 | 6.57 | 12.35 | 36.43 | 1.33 | 58.15 |
| **GLY485** | 7.51 | 5.29 | 8.75 | 4.58 | 6.49 | 7.60 | 51.89 | 26.92 | 9.20 |
| **PHE486** | 5.78 | 39.58 | 76.54 | 76.59 | 96.09 | 88.80 | 99.47 | 81.12 | 59.08 |
| **ASN487** | 47.18 | 90.23 | 99.29 | 99.29 | 94.67 | 94.00 | 77.65 | 79.43 | 69.52 |
| **CYS488** | 1.91 | 33.98 | 23.59 | 23.55 | 15.82 | 31.63 | 99.78 | 1.64 | 19.41 |
| **TYR489** | 94.09 | 99.16 | 100.00 | 99.91 | 99.96 | 99.07 | 100.00 | 84.45 | 99.78 |
| **PHE490** | 4.58 | 14.44 | 60.51 | 50.07 | 56.37 | 25.06 | 99.51 | 4.58 | 85.83 |
| **PRO491** | 0.93 | 0.13 | 0.27 | 0.00 | 2.75 | 0.00 | 10.93 | 49.62 | 98.22 |
| **LEU492** | 0.00 | 0.00 | 0.04 | 0.58 | 7.37 | 0.22 | 100.00 | 1.47 | 30.65 |
| **GLN493** | 69.79 | 81.52 | 98.76 | 99.91 | 98.49 | 99.91 | 100.00 | 79.43 | 99.16 |
| **SER494** | 22.97 | 29.76 | 90.23 | 32.47 | 2.98 | 90.36 | 100.00 | 43.71 | 99.69 |
| **TYR495** | 3.11 | 56.77 | 79.43 | 36.25 | 0.22 | 89.34 | 0.36 | 27.19 | 0.04 |
| **GLY496** | 47.13 | 57.71 | 54.82 | 34.92 | 40.47 | 86.85 | 87.69 | 0.62 | 12.39 |
| **GLN498** | 57.53 | 82.01 | 86.49 | 66.15 | 56.82 | 91.43 | 95.07 | 3.33 | 96.58 |
| **PRO499** | 19.55 | 0.44 | 0.00 | 0.00 | 0.00 | 0.09 | 19.19 | 3.55 | 59.71 |
| **THR500** | 12.13 | 69.08 | 3.33 | 16.75 | 5.55 | 53.89 | 2.93 | 0.00 | 69.70 |
| **ASN501** | 18.53 | 70.95 | 53.31 | 41.05 | 68.01 | 93.34 | 0.98 | 0.13 | 59.75 |
| **VAL503** | 3.20 | 88.32 | 7.24 | 0.04 | 0.18 | 0.00 | 0.00 | 0.00 | 10.17 |
| **GLY504** | 65.04 | 94.36 | 76.10 | 0.93 | 6.35 | 0.00 | 0.00 | 0.00 | 1.24 |
| **TYR505** | 97.07 | 99.60 | 99.96 | 37.05 | 58.02 | 46.56 | 0.62 | 0.13 | 48.11 |

^#^Delta: d1-3, Omicron o1-3, Wild type wt1-3.

Note: Data for all three sets of simulations are displayed separately.

Table S2B: Residue Wise contacts and their stabilities during the last 75% of the simulation time for the antibody interface.

| **Residue^$^** | **d1^#^** | **d2^#^** | **d3^#^** | **o1^#^** | **o2^#^** | **o3^#^** | **wt1^#^** | **wt2^#^** | **wt3^#^** |
| --- | --- | --- | --- | --- | --- | --- | --- | --- | --- |
| **GLU1H** | 0.22 | 0.00 | 0.00 | 2.22 | 30.92 | 4.31 | 1.55 | 1.60 | 63.35 |
| **VAL2H** | 0.76 | 0.00 | 0.00 | 7.24 | 36.25 | 1.87 | 2.27 | 2.49 | 56.95 |
| **SER25H** | 0.00 | 0.00 | 0.00 | 0.00 | 0.00 | 0.00 | 0.00 | 0.00 | 53.98 |
| **GLY26H** | 0.62 | 0.00 | 0.18 | 19.01 | 36.07 | 0.40 | 15.37 | 2.27 | 96.98 |
| **LEU27H** | 1.07 | 0.00 | 0.22 | 24.12 | 36.65 | 3.11 | 30.21 | 2.40 | 81.43 |
| **THR28H** | 10.26 | 10.71 | 4.66 | 32.65 | 35.36 | 4.40 | 97.73 | 14.22 | 96.09 |
| **SER30H** | 4.26 | 4.62 | 14.97 | 0.67 | 0.00 | 0.13 | 70.77 | 3.24 | 8.22 |
| **SER31H** | 43.94 | 81.52 | 91.47 | 53.44 | 35.98 | 31.19 | 100.00 | 18.66 | 99.78 |
| **ASN32H** | 28.08 | 34.92 | 46.38 | 14.17 | 32.56 | 34.83 | 78.54 | 16.35 | 96.53 |
| **TYR33H** | 61.97 | 86.63 | 96.49 | 89.29 | 71.83 | 41.40 | 100.00 | 50.42 | 99.91 |
| **TRP47H** | 0.00 | 21.59 | 57.22 | 0.18 | 0.80 | 16.79 | 47.18 | 35.63 | 0.00 |
| **TYR52H** | 54.95 | 75.74 | 89.78 | 97.29 | 87.61 | 54.33 | 100.00 | 94.49 | 98.40 |
| **SER53H** | 58.69 | 90.09 | 97.69 | 2.93 | 0.53 | 4.13 | 99.96 | 27.54 | 91.20 |
| **GLY54H** | 46.65 | 48.20 | 17.41 | 1.33 | 0.04 | 0.09 | 32.61 | 29.19 | 60.15 |
| **SER56H** | 40.29 | 29.94 | 79.65 | 58.64 | 33.36 | 21.86 | 48.11 | 71.92 | 73.39 |
| **THR57H** | 1.38 | 4.31 | 53.00 | 22.83 | 5.29 | 6.71 | 2.80 | 13.11 | 18.30 |
| **PHE58H** | 55.53 | 95.02 | 100.00 | 95.82 | 77.34 | 73.08 | 100.00 | 94.54 | 67.88 |
| **TYR59H** | 2.09 | 47.85 | 99.64 | 15.19 | 0.36 | 8.84 | 27.37 | 1.29 | 1.16 |
| **ASP61H** | 43.94 | 84.32 | 94.31 | 17.90 | 0.27 | 9.82 | 52.47 | 4.44 | 0.98 |
| **LYS64H** | 60.60 | 88.67 | 99.51 | 30.56 | 3.24 | 23.99 | 67.79 | 59.31 | 2.71 |
| **ARG97H** | 16.35 | 21.28 | 10.80 | 7.02 | 25.81 | 53.04 | 4.66 | 18.66 | 15.24 |
| **ASP98H** | 0.80 | 1.87 | 0.04 | 0.31 | 0.58 | 0.09 | 69.08 | 0.00 | 0.13 |
| **LEU99H** | 70.86 | 95.42 | 96.31 | 39.14 | 33.94 | 83.78 | 100.00 | 26.57 | 99.96 |
| **ASP100H** | 61.71 | 97.33 | 97.56 | 100.00 | 98.13 | 99.78 | 100.00 | 57.31 | 99.29 |
| **VAL101H** | 76.01 | 99.78 | 98.67 | 99.16 | 100.00 | 99.02 | 100.00 | 99.82 | 96.85 |
| **TYR102H** | 94.98 | 100.00 | 100.00 | 98.18 | 99.73 | 99.47 | 100.00 | 93.11 | 99.96 |
| **ASP1L** | 18.84 | 58.51 | 54.42 | 13.51 | 10.93 | 23.72 | 0.49 | 0.09 | 14.26 |
| **GLN27L** | 54.91 | 82.14 | 12.53 | 5.82 | 3.60 | 1.64 | 1.78 | 2.62 | 31.23 |
| **GLY28L** | 73.52 | 93.69 | 17.24 | 0.93 | 1.24 | 0.09 | 2.27 | 15.19 | 35.81 |
| **ILE29L** | 56.29 | 0.00 | 0.04 | 3.24 | 11.77 | 0.22 | 2.84 | 14.53 | 48.11 |
| **SER30L** | 43.49 | 99.91 | 94.18 | 2.04 | 36.74 | 19.90 | 5.78 | 61.97 | 27.72 |
| **SER31L** | 72.55 | 99.51 | 41.45 | 24.21 | 38.96 | 36.78 | 52.47 | 97.82 | 0.76 |
| **TYR32L** | 98.45 | 100.00 | 99.64 | 96.36 | 84.50 | 86.45 | 100.00 | 98.98 | 66.90 |
| **TYR49L** | 9.51 | 58.91 | 41.23 | 50.73 | 59.53 | 90.32 | 45.27 | 40.74 | 45.27 |
| **ALA50L** | 28.83 | 50.96 | 33.32 | 3.78 | 53.31 | 1.16 | 61.79 | 97.69 | 0.04 |
| **ALA51L** | 2.35 | 76.72 | 0.36 | 7.37 | 55.80 | 5.64 | 2.35 | 2.62 | 0.00 |
| **THR53L** | 11.64 | 12.48 | 12.17 | 44.78 | 55.84 | 59.00 | 29.72 | 44.60 | 29.81 |
| **LEU54L** | 0.09 | 0.40 | 0.00 | 28.61 | 33.05 | 77.08 | 0.00 | 0.31 | 5.02 |
| **GLN55L** | 0.00 | 2.00 | 0.09 | 28.65 | 2.18 | 88.98 | 0.04 | 0.18 | 3.29 |
| **SER56L** | 3.82 | 15.19 | 5.02 | 40.78 | 59.44 | 87.29 | 0.18 | 3.02 | 31.27 |
| **SER67L** | 3.51 | 72.46 | 1.42 | 0.00 | 0.04 | 0.00 | 0.00 | 13.42 | 0.00 |
| **ASN92L** | 80.54 | 99.42 | 77.92 | 96.93 | 98.49 | 82.32 | 100.00 | 76.50 | 75.21 |
| **SER93L** | 15.24 | 67.44 | 0.13 | 25.94 | 41.58 | 5.20 | 87.74 | 0.31 | 54.42 |
| **TYR94L** | 99.87 | 73.39 | 100.00 | 99.82 | 85.43 | 99.42 | 100.00 | 99.87 | 84.67 |
| **PRO95L** | 62.95 | 91.12 | 99.78 | 89.12 | 45.09 | 93.07 | 98.13 | 85.92 | 7.11 |
| **PRO96L** | 15.90 | 10.93 | 3.38 | 0.84 | 8.84 | 8.04 | 99.91 | 51.31 | 0.84 |
| **LYS97L** | 11.68 | 2.09 | 71.57 | 59.75 | 65.75 | 71.57 | 100.00 | 68.50 | 69.52 |

^#^Delta: d1-3, Omicron o1-3, Wild type wt1-3.

^$^L and H are indicating the antibody chain identifier.

Note: Data for all three sets of simulations are displayed separately.

Table S3: Average end state free energies (MM-GBSA) for all the variants during three independent simulation runs using the most populated structure.

| **Simulation run** | **WT^1^** | **Delta^1^** | **Omicron^1^** |
| --- | --- | --- | --- |
| I | -193.68 | -85.60 | -60.88 |
| II | -90.01 | -79.54 | -55.19 |
| III | -131.24 | -117.54 | -51.11 |
| Average | -138.31 ± 51.2 | -94.23 ± 20.42 | -55.73 ± 4.91 |

^1^Energy values are reported in kcal/mol along with the standard deviation.
